# A Mathematical Framework for Comparison of Intermittent versus Continuous Adaptive Chemotherapy Dosing in Cancer

**DOI:** 10.1101/2024.02.19.580916

**Authors:** Cordelia McGehee, Yoichiro Mori

## Abstract

Chemotherapy resistance in cancer remains a barrier to curative therapy in advanced disease. Dosing of chemotherapy is often chosen based on the maximum tolerated dosing principle; drugs that are more toxic to normal tissue are typically given in on-off cycles, whereas those with little toxicity are dosed daily. When intra-tumoral cell-cell competition between sensitive and resistant cells drives chemotherapy resistance development, it has been proposed that adaptive chemotherapy dosing regimens, whereby a drug is given intermittently at a fixed dose or continuously at a variable dose based on tumor size, may lengthen progression free survival over traditional dosing [1]. Indeed, in mathematical models using modified Lotka Volterra systems to study dose timing, rapid competitive release of the resistant population and tumor outgrowth is apparent when cytotoxic chemotherapy is maximally dosed [2]. This effect is ameliorated with continuous (dose modulation) or intermittent (dose skipping) adaptive therapy in mathematical models and experimentally, however, direct comparison between these two modalities has been limited. Here, we develop a mathematical framework to formally analyze intermittent adaptive therapy in the context of bang-bang control theory. We prove that continuous adaptive therapy is superior to intermittent adaptive therapy in its robustness to uncertainty in initial conditions, time to disease progression, and cumulative toxicity. We additionally show that under certain conditions, resistant population extinction is possible under adaptive therapy or fixed-dose continuous therapy. Here, continuous fixed-dose therapy is more robust to uncertainty in initial conditions than adaptive therapy, suggesting an advantage of traditional dosing paradigms.

**Significance Statement:** Intra-tumoral cell-cell competition between drug resistant and drug sensitive cancer cells is hypothesized to drive resistance development after chemotherapy administration in certain cancers. Optimization of chemotherapy delivery through dose or schedule adaptation based on tumor parameters (adaptive therapy) has been hypothesized to utilize chemotherapy sensitive cells to compete with chemotherapy resistant cells and delay tumor outgrowth. Here, we develop an analytical framework to compare two alternative adaptive therapy approaches: dose modulation (continuous) and schedule modulation (intermittent). Through direct analytical comparison, we demonstrate that continuous adaptive therapy is superior to intermittent adaptive therapy across physiologically relevant metrics. Furthermore, in parameters spaces where resistant population extinction is feasible, traditional fixed-dose therapy at a low dose may carry additional benefits over adaptive therapy.

## Introduction

Thanks to advances in medicine and cancer treatments, more and more patients are living longer with metastatic cancer [3, 4]. In these patients, a cure is often not possible and the goal is to limit overall side effects from cancer treatments, prolong progression free survival, and limit symptoms from the cancer itself [5].

Chemotherapy dosing has traditionally derived from the *maximum tolerated dose* (MTD) principle [6] (Figure 1). For chemotherapy or combination chemotherapy that is associated with high levels of toxicity for patients, dosing is often based on how often the chemotherapy can be given without incurring severe side effects as measured in Phase 2 clinical trials. The final dosing schedule is typically fixed (every 2 weeks, every 3 weeks, etc) as is the number of doses that a patient ultimately receives (termed “cycles”). The dosing schedule can be adjusted based on patient status (e.g. dosing may be delayed if a patient becomes sick) or discontinued if the patient’s tumor is not responding to treatment. As an alternative to MTD, *metronomic dosing*, where a chemotherapy is administered at a lower dose and given at regular intervals, has been utilized to minimize toxicity and control disease progression [7, 8]. For certain chemotherapeutic agents that are relatively non-toxic and available in oral formulation, dosing may be given daily as tolerated, leading to a stable steady state concentration of drug in the body (continuous therapy). This type of treatment can be considered metronomic therapy and is used alone or in combination with other chemotherapeutic agents and dosing strategies [9]. Ultimately, many advanced metastatic cancers progress through standard treatment and become refractory to chemotherapy.

**Figure 1:**
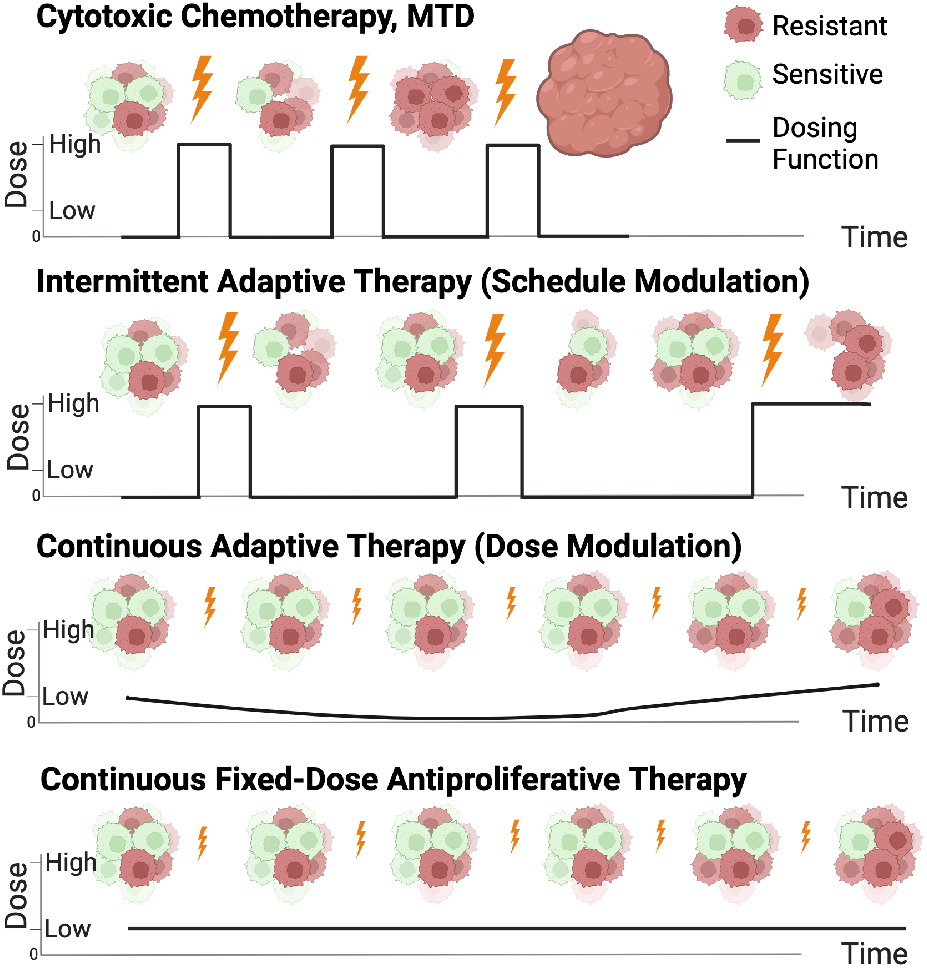
Illustration comparing four distinct dosing paradigms in cell-cell competition models in cancer. Cytotoxic chemotherapy at maximum tolerated dose (MTD) results in early extinction of the sensitive cell population. Two adaptive therapy models are shown: intermittent adaptive therapy (resulting from on-off dose modulation aka “dose-skipping”) and continuous adaptive therapy (resulting from dose modulation to maintain a given tumor volume). Finally, fixed low dose therapy in the antiproliferative range of drug action is shown and is considered a type of metronomic therapy. *Created with BioRender.com*.

In broad terms, a tumor is defined as sensitive to a given chemotherapy if it shrinks or does not grow with treatment. A tumor is defined as resistant to chemotherapy if it continues to grow in the presence of treatment. While these terms are applied to entire tumors, in reality, tumors are heterogeneous and may contain subpopulations of drug resistant and sensitive cells. Pancancer analysis of intratumoral tumor heterogeneity has suggested that high heterogeneity is correlated with decreased overall survival across a variety of cancer types [10]. The mechanisms by which interactions between these subpopulations contribute to treatment response and resistance is likely highly dependent on the type of cancer, the tumor microenvironment and architecture, and the resistance mechanism in question.

Adaptive chemotherapy administration, wherein a drug schedule or dose is dependent on tumor size (or some other metric of tumor burden), has been proposed as a method to lengthen progression free survival in cancers where a resistant subpopulation already present in the tumor is likely to drive disease progression [1, 11, 2]. This method has shown some success in increasing progression free survival in a clinical trial of patients with advanced metastatic prostate cancer [2] and in preclinical models of ovarian cancer, breast cancer, and melanoma [1, 12, 13]. There are planned and ongoing clinical trials to evaluate adaptive strategies in the treatment of rhabdomyosarcoma, prostate cancer, melanoma, ovarian cancer, and advanced basal cell carcinoma [14].

The rationale for implementation of adaptive therapy is based largely on two ecological principles: fitness cost and competitive release. In the context of cancer cell competition, fitness cost is the idea that becoming chemotherapy resistant inherently leads to a trade-off in some other area of ecological fitness [1]. The observation that slower growth rates of cancer cell subpopulations can lead to chemoresistance to traditional cytotoxic chemotherapy [15] can be used as evidence of this cost. Additionally, resistant cell upregulation of drug efflux pumps which consume large amounts of cellular ATP [16] is a plausible mechanism by which a fitness cost to resistance is incurred for certain drugs. However, whether or not fitness cost in the context of slower growth of chemoresistant cells or other mechanisms applies more widely across newer targeted inhibitor therapies remains to be seen. Recent exploration of the adaptive therapy models suggests that even without significant differences in growth rate between sensitive and resistant cells, there is still a progression free survival advantage with adaptive therapy over traditional fixed dosing schemes [17].

The observation of continued efficacy of adaptive chemotherapy even without a fitness cost of resistance derives from the ecological principle of competitive release. Assuming individual cancer cells are competing for resources (space, blood flow, nutrients, etc), then the ability of cells to proliferate is dependent upon the overall density of cells. Thus, the closer a tumor is to its *carrying capacity* (the number of tumor cells that can be sustained by the local environment), the lower the growth rate of the tumor. In traditional dosing schemes, the goal of treatment is to maximally shrink the tumor which results in rapid reduction of the drug sensitive species and thus tumor volume. With this reduction in overall tumor volume, the resistant cells are able to proliferate faster. The observed rapid proliferation of the resistant cells is termed *competitive release*. Under adaptive dosing strategies, however, the sensitive cell population is purposefully maintained within the tumor in order to inhibit competitive release of the resistant cell population [1, 2, 11].

Adaptive therapy strategies can be broadly categorized as intermittent or continuous. In *intermittent adaptive therapy*, a fixed dose of drug is given until a tumor reaches a certain lower size threshold and then withdrawn until the tumor regrows to a certain upper size threshold (Figure 1). This has also been coined “on-off” adaptive therapy or “dose-skipping” adaptive therapy [14]. In *continuous adaptive therapy*, on the other hand, a drug is given continuously with variable dosing to hold a tumor to a given volume, also termed “dose modulation” adaptive therapy [14] (Figure 1). Direct comparison between intermittent adaptive therapy and continuous adaptive therapy has been explored experimentally [12] and computationally [18] in an agent based model. Mathematically, the observation that higher tumor volume implies slower growth has demonstrated that continuous adaptive therapy held at the highest tolerated tumor volume maximizes time to treatment failure and may be superior to intermittent adaptive therapy [19]. However, formal mathematical analysis of intermittent adaptive therapy has been limited due to the discontinuous nature of the dosing function.

In this work, we show that intermittent adaptive therapy can be mathematically modeled as a bang-bang control problem [20] and that, under certain conditions, the resistant population can be driven to extinction. We prove that the analytical limit as the upper and lower control boundaries of adaptive therapy approach each other yields a continuous dosing function and that this continuous dosing function is equivalent to continuous adaptive therapy. Using this result, we show that continuous adaptive therapy is ideal to intermittent adaptive therapy in time to resistant subpopulation outgrowth. In the case where the resistant population can be driven to extinction, we show that when compared to intermittent adaptive therapy, continuous adaptive therapy is more robust to uncertainty in initial conditions, yields a quicker time to resistant population eradication, and carries the lowest cumulative toxicity. Finally, we show that the continuous dosing function is in the antiproliferative range of drug action and thus can be directly compared with fixed low-dose continuous therapy, a type of metronomic therapy (Figure 1). An unexpected advantage of this continuous metronomic therapy over adaptive therapy is that it can result in resistant extinction under an often broader set of initial conditions than adaptive therapy. Furthermore, fixed-dose therapy is biologically and clinically easier to implement. To our knowledge, this is the first direct analytical comparison between continuous and intermittent adaptive therapy dosing regimes in cancer and demonstrates the utility of this benchmarking approach to compare continuous control models with bang-bang control models for understanding tumor evolution under drug perturbation.

## Results

### Mathematical Model

The Lotka Volterra equations are a well established dynamical system for studying competition between species [21]. For the purposes of this manuscript, the two species Lotka Volterra model is described below for the sensitive cell population, *x*, and the resistant cell population, *y*, under logistic growth. A conceptual representation of this model is shown in Figure 2A.

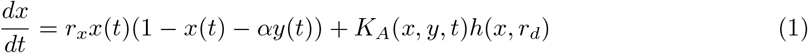

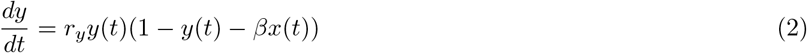

The major difference between this model and a traditional competition model is the presence of an indicator function *K*_*A*_(*x, y, t*) in Equation 1 which takes values 1 or 0 depending on whether the drug is on or off respectively. The effect of the drug on the sensitive cell line is given by the function *h*(*x, r*_*d*_) where *r*_*d*_ is the “dose” of drug. Note that the carrying capacity for each species is assumed to be equal and is normalized to 1.

**Figure 2:**
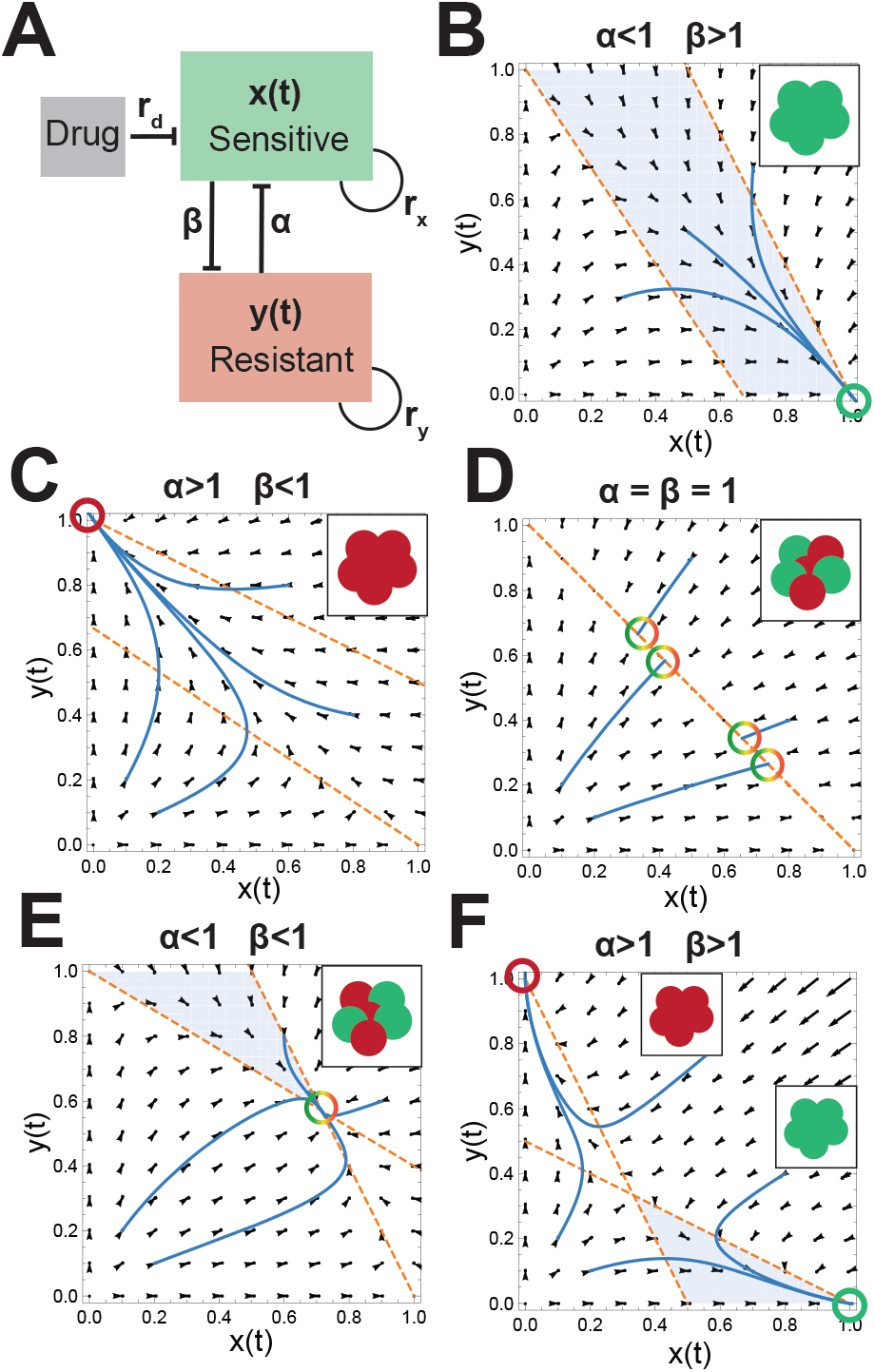
Cell-cell competition model across parameter spaces. **(A)** Conceptual representation of cell-cell competition model. The growth rates of the sensitive and resistant populations are given by *r*_*x*_ and *r*_*y*_ respectively. *α* and *(3* are the competition parameters of *y* on *x* and *x* on *y* respectively. Drug dose is given by *r*_*d*_ and acts only to decrease the sensitive species. **(B-F)** Vector fields corresponding to different relative competition strengths between the sensitive and resistant cell populations without drug. The nullclines are plotted in orange. The equilibrium states are shown as illustrations in the upper right insets and areas where *x*′ > 0 and *y*′ < 0 are shaded blue.

Drug effect in this paper is modeled as removal of the sensitive cell population and is assumed to be of the form below:

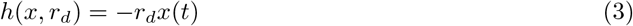

Representing the drug action in this manner allows for a range of drug effect from *antiproliferative* (decreasing the growth rate of the sensitive cells) to *cytotoxic* (leading to sensitive cell extinction) based on administered dose. (See Supplementary Figure 8 and Supplementary Section 1.3.)

The behavior of this model in the absence of drug is shown in Figure 2B-F. Note that when the sensitive or resistant cell line is a strong competitor (*α* or *β* > 1 respectively) relative to weak competition (*β* or *α* < 1) from the other subpopulation, the equilibrium state is composed of the single stronger species at carrying capacity while the weaker subpopulation vanishes (Figure 2B and 2C). On the other hand, when both are weak competitors (2E) an equilibrium heterogeneous state exists, and when both are strong competitors (2F) the tumors trend to homogeneous populations determined by the initial conditions. Finally, if competition terms are both equal to 1, the heterogeneous tumor is stable with the subpopulation ratio dependent on initial conditions. Note that the growth rate (given by *r*_*x*_ and *r*_*y*_) for each species does not affect the position of the nullclines in this model.

Importantly, if the goal of treatment is to reduce the resistant population to near 0, then the natural strategy is not to treat in parameter spaces in which the system equilibrium at carrying capacity results in an entirely sensitive tumor. Because of this, we assume that the maximal tumor size (sensitive plus resistant cells) must be maintained at a volume less than carrying capacity and we call this the *upper control line, A*.

### Intermittent Adaptive Therapy

Intermittent adaptive therapy is modeled using bang-bang control where the controller is turned on (*K*_*A*_ = 1) when the total tumor volume (resistant + sensitive) reaches the upper control line, *x* + *y* = *A*. Once the tumor volume has decreased to the lower control line *A - δ*, the controller is then turned off (*K*_*A*_ = 0) and the tumor regrows. The dose of drug is held constant at *r*_*d*_ which is assumed to be in the cytotoxic range. This strategy is continued indefinitely or until the tumor is no longer controllable (e.g. the tumor size cannot be decreased to *A - δ* with administration of drug).

In the case where the resistant cell population is a weaker competitor than the sensitive cell population (*α* < 1 and *β* > 1, Figure 2B), the relative position between the upper and lower control lines and the *y*-nullcline yields the possibility of three distinct behaviors that are dependent on the initial tumor composition. An example set of trajectories and phase space is shown in Figure 3A for varying initial conditions demonstrating sensitive cell population extinction (Figure 3B) sensitive/resistant equilibrium (Figure 3C), and extinction of the resistant cell population (Figure 3D).

**Figure 3:**
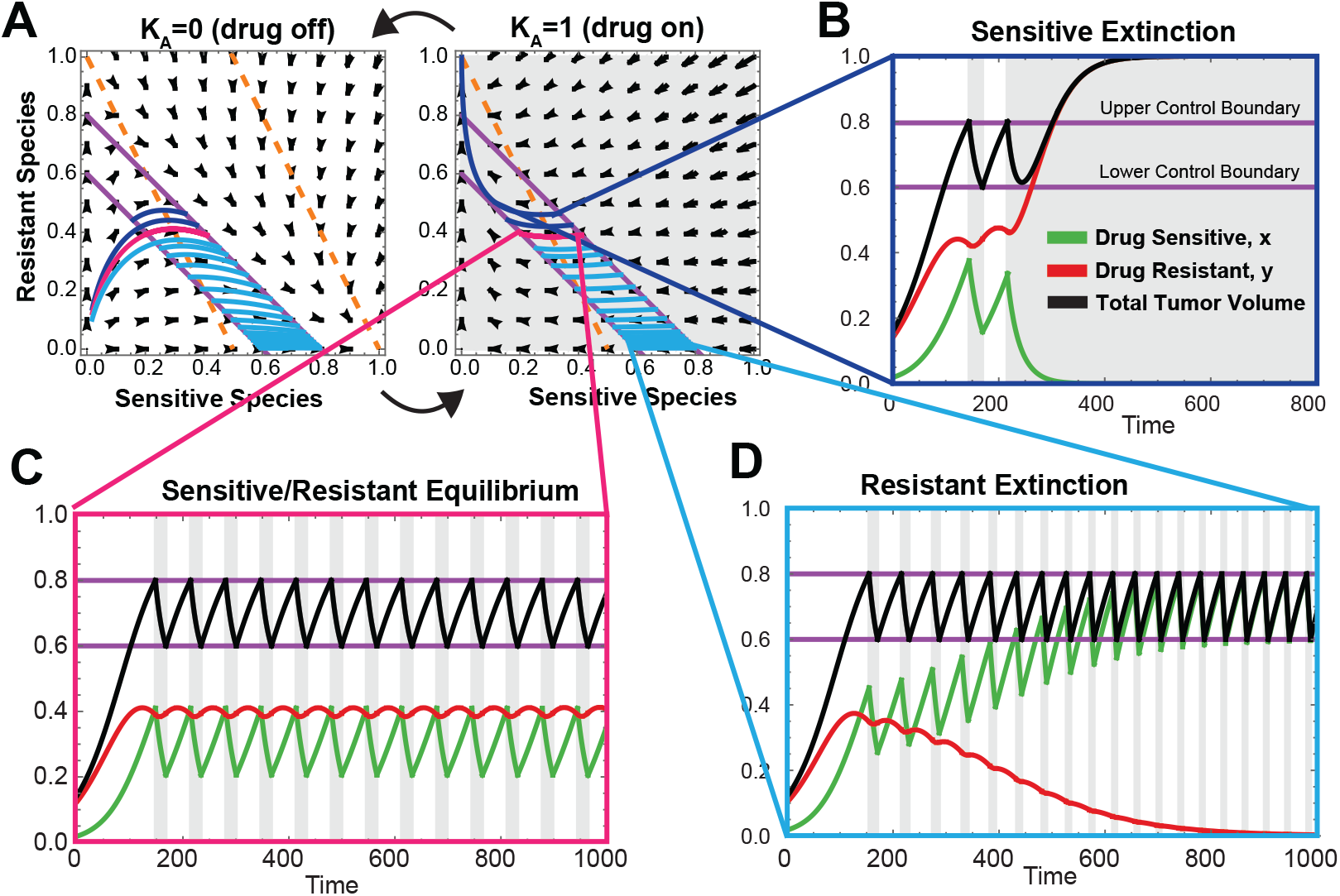
Simulation of intermittent adaptive therapy for *α* < 1, *β* > 1 showing three possible outcomes. (A) Phase space for drug on and drug off with 3 trajectories of adaptive therapy plotted (light blue, pink, and dark blue) under the same simulation parameters with different initial conditions. (B) Sensitive extinction time course corresponding to the dark blue trajectory in phase space shown in A. (C) Sensitive/resistant equilibrium time course corresponding to the pink trajectory in phase plane shown in A. (D) Resistant extinction time course corresponding to the light blue trajectory in phase space shown in A.

The case where the resistant cell population is the stronger competitor yields only one outcome (sensitive extinction) and is shown in Supplemental Figure 10 and discussed in Supplemental Section 2.1.2. In this case, while resistant extinction is not achievable, adaptive therapy lengthens time to tumor progression over continuous treatment with the same dose of drug or no treatment which has been previously reported [17].

### Optimal Intermittent Adaptive Therapy Limits to Continuous Adaptive Therapy

Given that a higher maintained sensitive cell population results in lower resistant cell population growth rate, decreasing *δ* with fixed upper tumor volume intuitively increases the time to resistant outgrowth under conditions leading to sensitive cell population extinction and decreases time to resistant extinction under conditions leading to resistant cell population extinction. With this intuition in mind, we formally explore the analytical limit of adaptive therapy as *δ →* 0 for fixed upper control line *x* + *y* = *A*. See Supplemental Sections 2.2 - 2.5 for mathematical discussion.

We prove that intermittent adaptive therapy converges to the *x* + *y* = *A* line in this limit (Supplemental Theorem 2.7) and formally define a control region in phase space where the limiting behavior converges (Supplemental Section 2.3). We similarly define the strict control region to be the portion of phase space where the the limit as *δ →* 0 of intermittent adaptive therapy is always in control and the tumor volume never increases above *A* (Supplemental Definition 2.3).

A simulation of this limiting system is shown in Figure 4C. Note that the limiting trajectory at *δ →* 0 recapitulates the behavior of the oscillating system shown in Figure 3. At the point of intersection between the control line and the y-nullcline, an unstable equilibrium exists corresponding to the equilibrium seen in Figure 3B. Similarly, resistant extinction and sensitive extinction are determined based on the intersection of the initial trajectory with the *x* + *y* = *A* line.

**Figure 4:**
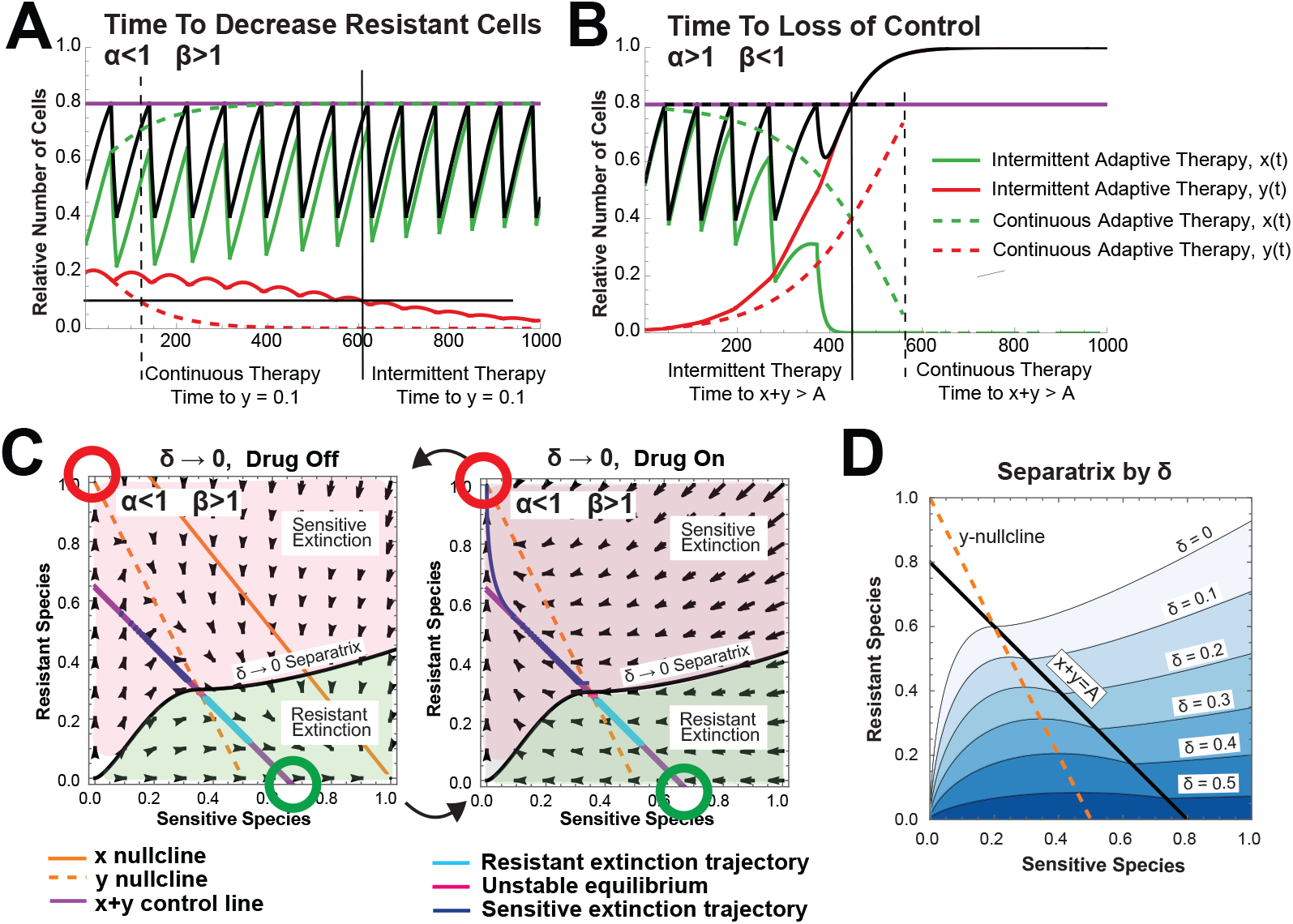
Time to disease progression, time to loss of control, and robustness to uncertainty in initial conditions are optimized under continuous adaptive therapy. (**A**) Time to decrease resistant cell population to threshold (*y* = 0.01) when *α* < 1 and *β* > 1 under intermittent adaptive therapy (solid lines) and continuous adaptive therapy (dotted lines). (Model Parameters: *α* = 0.5, *β* = 2, *r*_*d*_ = 0.1, *r*_*x*_ = 0.03, *r*_*y*_ = 0.02) (**B**) Time to loss of control when *α* > 1 and *β* < 1 under intermittent adaptive therapy (solid lines) and continuous adaptive therapy (dotted lines). (Model Parameters: *α* = 1.5, *β* = 0.7, *r*_*d*_ = 0.1, *r*_*x*_ = 0.03, *r*_*y*_ = 0.02) (**C**) Phase planes corresponding to < *→* 0 trajectories with the numerically computed separatrix delineating between initial conditions leading to resistant species extinction and those leading to sensitive species extinction. (**D**) Numerically computed separatrix for varying values of < (Model Parameters: *α* = 0.5, *β* = 2, *r*_*d*_ = 0.05, *r*_*x*_ = 0.03, *r*_*y*_ = 0.02). Theoretical computation of the separatrix is discussed in Supplemental Section 2.4 and has excellent concordance with numerical results.

The observation that intermittent adaptive therapy converges in the limit as *δ →* 0 can be used to compute a continuous dosing function as a function of the sensitive cell population (Supplemental Equation 30). This continuous dosing function is a smooth function essentially representing a time averaged *r*_*d*_ and is equivalent to continuous adaptive therapy on the *x* + *y* = *A* line by definition. The shape of the dosing function, 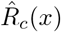, depends on the underlying phase space. In certain cases, 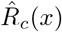 represents a decreasing function consistent with reported experimental observations [12] (Supplemental Figure 18). When resistant extinction can be achieved, this function takes values in the anti-proliferative range of drug action (Supplemental Corollary 2.6). Interestingly, the dosing function 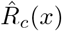 does not depend on the original intermittent adaptive therapy dose, suggesting that it represents an absolute theoretical limit independent of *r*_*d*_.

The knowledge that intermittent adaptive therapy limits to continuous adaptive therapy as *δ* decreases allows for formal comparison of these two adaptive therapy modalities. First, we show that the time to loss of control under continuous adaptive therapy is maximized relative to the time to loss of control under intermittent adaptive therapy (Supplemental Section 2.6). Similarly, the time to resistant extinction (when possible) is minimized under continuous adaptive therapy (Supplemental Section 2.6 and Figure 14A). An example simulation of these results is shown in Figure 4A and 4B. While the limit of time to disease control or loss of control as *δ →* 0 converges to the computed time in continuous adaptive therapy, the same is not true of drug toxicity. Specifically, we show that when cumulative drug toxicity is modeled as an increasing nonlinear function of drug dose, then optimal drug toxicity as *δ →* 0 is strictly larger under intermittent adaptive therapy than continuous adaptive therapy (Supplemental Section 2.7 and Figure 14B). This suggests an absolute advantage of continuous adaptive therapy over optimal intermittent adaptive therapy.

Finally, in addition to drug toxicity, one of the most important aspects of a therapeutic strategy is robustness to uncertainty in parameters. Specifically, the relative composition of a tumor in terms of sensitive versus resistant subpopulations may be difficult, if not impossible, to attain precisely. We note that as *δ* decreases, the parameter space of tumor compositions that lead to resistant population extinction increases (See Figure 4D). Mathematically, it is shown that the strict control region for continuous adaptive therapy is strictly greater than the strict control region for intermittent adaptive therapy (See Supplemental Section 2.4). This suggests a further advantage of continuous adaptive therapy over intermittent adaptive therapy in cases where resistant extinction is possible.

### Continuous Fixed-Dose Treatment is Non-Inferior to Adaptive Therapy For Resistant Population Extinction

In the previous section, we showed that in regions where resistant subpopulation extinction is possible, continuous adaptive therapy is superior to intermittent adaptive therapy in minimization of the time to resistant population extinction, minimization of overall drug toxicity, and maximization of initial conditions leading to resistant subpopulation extinction. As resistant subpopulation extinction yields the possibility of total tumor eradication, it presents an enticing area for further analysis. Accordingly, we sought to further characterize this region to understand the relative benefits and limitations of adaptive therapy versus standard fixed-dose therapies.

Analysis of this model under continuous fixed-dose drug administration again demonstrates that continuous cytotoxic doses of chemotherapy lead to rapid release of resistant cells and tumor outgrowth (Figure 5B). Antiproliferative drug doses (*r*_*d*_ < *r*_*x*_), on the other hand, act to decrease the sensitive species carrying capacity and slow their growth rate in the absence of a competing resistant cell line (Figure 5A and C and Supplemental Section 1.3). Similar to the results shown for adaptive therapy, fixed doses of antiproliferative drug can be identified that maintain the sensitive cell population indefinitely and lead to extinction of the resistant cell population. Interestingly, when a weakly competitive resistant cell population competes with the sensitive one, the same anti-proliferative dose of drug that results in sensitive cell population maintenance in the absence of competition may become “effectively cytotoxic” and trigger extinction of the sensitive cell population (Figure 5B, Supplemental Figure 16, and Supplemental Section 3.1). This result has implications beyond adaptive therapy. Specifically, drugs that are measured to be anti-proliferative *in vitro* or under conditions where nutrients and space are not growth limiting may be effectively cytotoxic in more realistic tumor environments where growth is constrained by the surrounding tissue and a competing normal cell population. Finally, in the case where the resistant cells are strong competitors, resistant cell population extinction is not achievable under continuous fixed dose therapy. Here, lower doses of drug result in increasing times to resistant cell population outgrowth (Supplemental Section 3.1).

**Figure 5:**
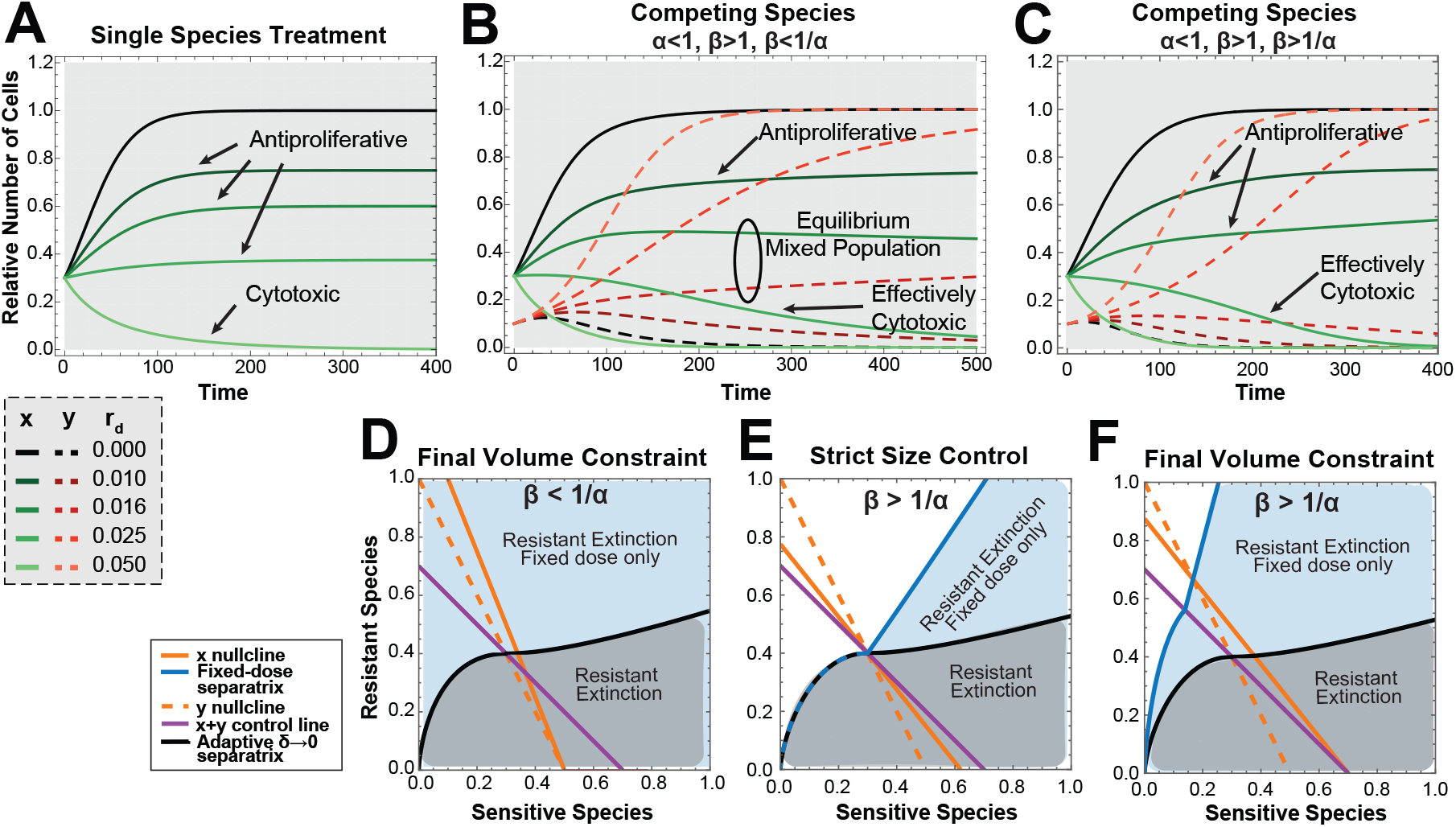
Resistant population extinction under continuous fixed dose treatment versus optimal adaptive therapy. (**A**) Treatment of the single species sensitive cell population alone versus (**B**) treatment with the same doses of drug (*r*_*d*_) in a mix of sensitive and resistant competing cells when *β* < 1*/α* and (**C**) when *β* > 1*/α*. (**D**) Phase plane when 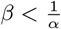 with the continuous adaptive < = 0 separatrix shown in black with initial conditions under the line leading to resistant extinction (shaded grey). All initial conditions lead to resistant extinction under fixed-dose continuous therapy (shaded blue) with the minimum achievable tumor volume taken when *r*_*d*_ is set such that the intersection of the x and y nullclines occurs on the x-axis. (**E**) Phase planes when 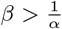 with the continuous adaptive < = 0 separatrix shown in black with initial conditions under the line leading to resistant extinction (shaded grey). In the first phase plane, *r*_*d*_ is chosen such that for all initial conditions where continuous adaptive control leads to resistant population extinction, continuous fixed-dose treatment will also yield resistant species extinction with final tumor size smaller than *A*. (**F**) In the second phase plane, *r*_*d*_ is set such that the equilibrium under fixed-dose control for favorable initial conditions (shaded blue) is set to *A. Simulation Parameters:* **A-C**, *r*_*x*_ = 0.04, *r*_*y*_ = 0.03, (*x*_0_, *y*_0_) = (0.3, 0.1) except A where *y*_0_ = 0. In B, *α* = 0.5 and *β* = 1.5. In C, *α* = 0.9 and *β* = 2.

In cases where resistant subpopulation extinction is achievable under both continuous fixed dose therapy and continuous adaptive therapy, we analyzed the minimum tumor volume at which the tumor could be maintained. This is divided into two cases. When *β* < 1*/α*, this minimum volume is explicitly solvable (see Supplementary Section 3.2) and is shown to be the minimum tumor volume by phase plane argument. Knowing this minimum tumor volume allows us to calculate the drug dose *r*_*d*_ needed to achieve control at the minimum achievable volume (See Supplementary Proposition 3.1) under any initial tumor composition. Continuous adaptive therapy, on the other hand, results in resistant species extinction only under certain initial conditions. Therefore, it is less robust to uncertainty in initial conditions than continuous fixed dose therapy (Figure 5C). It is important to note that in the regions where continuous fixed dose therapy can result in resistant population extinction and adaptive therapy cannot, the tumor size must increase above the upper tumor control volume, A, for a period of time even though steady state maintenance of tumor volume will be at or below A. The optimal strategy to minimize the size of tumor outgrowth above A is to wait to initiate continuous fixed dose treatment until the tumor volume reaches A.

For the second case when *β* > 1*/α*, a different minimum achievable tumor volume is given under continuous fixed-dose therapy and there is a dependence on initial conditions. However, the space of initial conditions leading to resistant extinction is always greater than or equal to that in continuous adaptive therapy for the same final tumor volume constraint (Figure 5D and Supplemental Section 3.3). Furthermore, there are a set of initial conditions under which resistant species extinction can be achieved under fixed-dose therapy but not adaptive therapy. Again, the tumor volume in these cases may increase above *A* transiently and this increase can be minimized by initiating treatment when the tumor size reaches *A*.

In summary, direct comparison of continuous adaptive therapy versus continuous fixed-dose therapy demonstrates that both strategies may lead to resistant extinction under certain conditions and that the set of initial conditions leading to resistant species extinction under fixed-dose therapy is broader than those under adaptive therapy. In cases where resistant control is not possible under adaptive therapy but is possible with continuous fixed dose therapy, the tumor size will exceed the upper control line for a period of time to achieve resistant species extinction. This effect is minimized if the treatment is initiated once tumor volume reaches the upper control line. Taken together, we conclude that continuous fixed-dose therapy is mathematically non-inferior to continuous adaptive therapy in achieving resistant species extinction and that if tumor volume need not be strictly controlled, then continuous fixed-dose therapy may be superior. Given the practical considerations of ease of dosing for fixed-dose therapy, this strategy may be preferable in practice.

## Discussion

In this study, a modified Lotka-Volterra model of adaptive therapy is described that can be used to analytically examine intermittent adaptive therapy as a bang-bang control problem. It is shown that the limit of this intermittent dosing as the lower dosing threshold approaches the upper control line converges to the upper control line and has equivalent behavior to continuous adaptive therapy. In cases where resistant population control is not achievable, continuous adaptive therapy is optimal to intermittent cytotoxic dosing in delaying resistant cell population outgrowth. These results are consistent with results put forward by Viossat *et. al*. [19] but differ in their theoretical connotation. Specifically, it is proven that intermittent adaptive therapy is equivalent to continuous adaptive therapy in the limit irrespective of the given fixed intermittent dose and that continuous adaptive therapy provides an upper boundary to idealized intermittent adaptive therapy behavior. We further prove that cumulative drug toxicity is strictly lower in continuous adaptive therapy over intermittent adaptive therapy, suggesting an absolute clinical advantage of this treatment modality. We additionally define a set of initial conditions for which resistant population extinction is achievable when the sensitive cell line is a stronger competitor than the resistant cell line (“fitness cost”). We show that for these conditions, continuous adaptive therapy is superior to intermittent adaptive therapy in terms of the time to resistant population extinction, cumulative drug toxicity, and robustness to uncertainty in initial conditions (Figure 6). While adaptive therapy clearly outperforms cytotoxic continuous fixed-dose therapy, continuous antiproliferative fixed-dose therapy, a type of metronomic therapy, offers increased robustness to uncertainty in initial conditions over adaptive therapy provided that tumor size need not be strictly contained for the entirety of treatment (Figure 6). We therefore conclude that in this region, continuous fixed-dose therapy is mathematically non-inferior to continuous adaptive therapy. As metronomic dosing is well established in practice and clinical requires less frequent monitoring, the practical considerations likely favor this therapy over continuous adaptive therapy.

**Figure 6:**
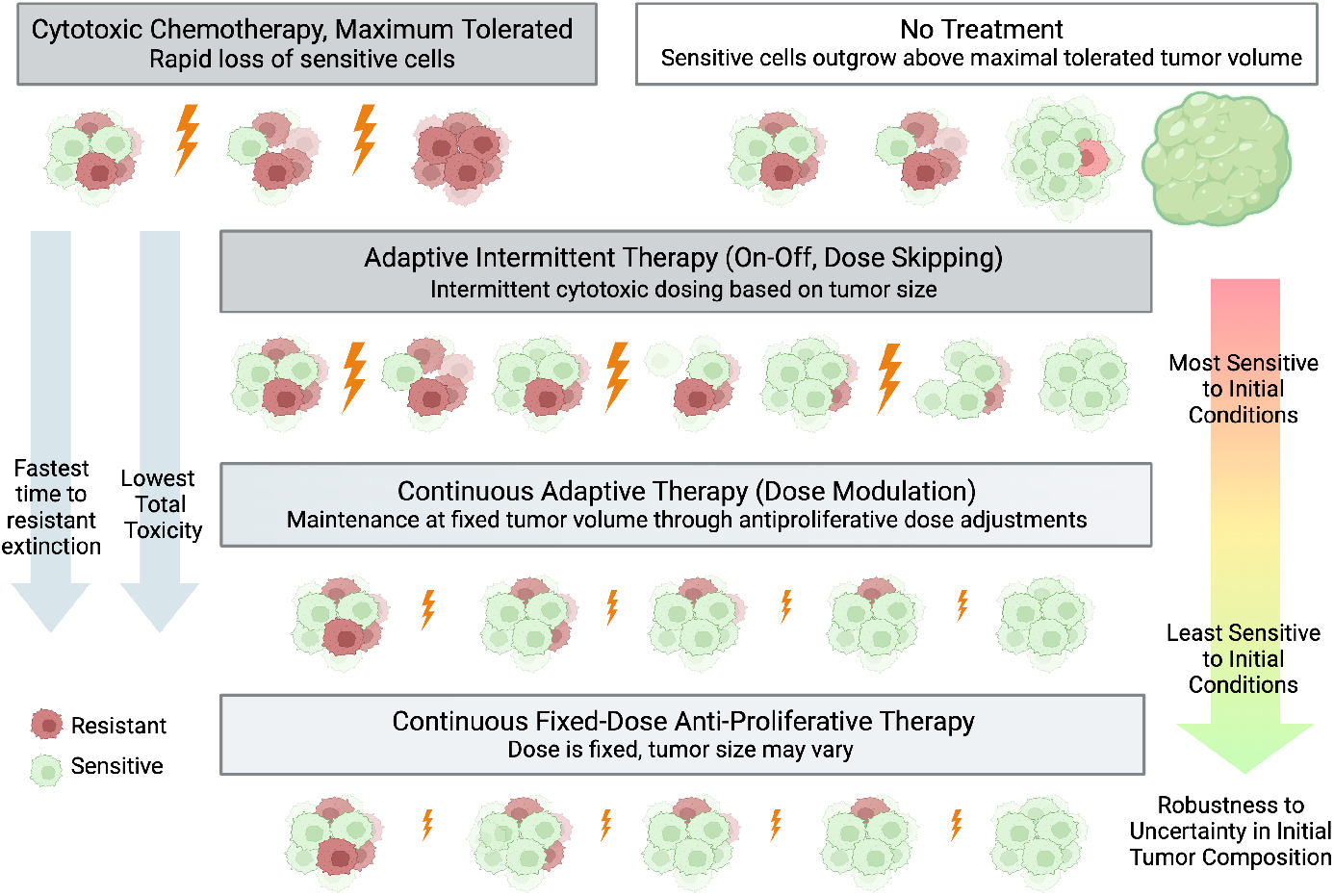
Summary of mathematical conclusions under conditions where resistant extinction is possible. *Created with BioRender.com*

Exploration of the space where resistant species extinction is possible has been previously limited as it necessitates strong competition from the sensitive cell line and a maintenance at a tumor volume that is high relative to the overall carrying capacity [19]. However, it is possible that combination therapy targeted at weakening the competitiveness of the resistant cell line and/or modulating the carrying capacity of the space may create circumstances where therapy aimed at extinction of the resistant cell population could be employed. As extinction of the resistant cell population creates the opportunity for tumor eradication at the end of therapy, it represents an important and enticing area for formal analysis.

This study is intended to provide a conceptual mathematical framework for comparing intermittent and continuous adaptive therapy against standard dosing regimes including MTD and continuous metronomic therapy in cases where competition between sensitive and resistant cell population drive resistance development. The presented work is purely mathematical in nature and limited by a lack of *in vitro* or *in vivo* data to validate the model of drug action, cell competition, and logistic growth. This model does not take into account spatial heterogeneity and assumes that populations of cells are well mixed. Additionally, this model does not account for accumulation of mutations through the tumor growth cycles that could occur in adaptive therapy.

In addition to the limitations of this particular model, it is important to note that the implementation of adaptive therapy itself is limited by biological and technical considerations. For example, if a resistant subpopulation of cells does not exist within a tumor, there is no benefit to adaptive therapy and the implementation of this strategy may lead to the accumulation of additional mutations as well as potential clinical morbidity. These strategies are additionally limited by the ability to accurately track tumor volume in time. Initial clinical trials have been performed in prostate cancer with the biomarker PSA as a proxy for tumor volume [2], however, it is unclear how best to track tumor volume in other tumor types where biomarkers corresponding to tumor size are not readily available. Tracking tumor volumes additionally makes the administration of continuous adaptive therapy potentially more difficult than intermittent adaptive therapy due to the need for real-time tumor volume tracking. These considerations are ameliorated somewhat by continuous fixed-dose therapy, however, choosing the correct initial dose depends on knowledge of tumor parameters which may not be feasible to attain. These considerations have been addressed in recent work aimed at drug titration algorithms to determine a stabilization doses of chemotherapy [22, 19, 23].

Despite the above limitations, several unexpected theoretical advantages of adaptive therapy exist including the observation by Enriquez-Navas *et. al*. that adaptive therapy schemes may maintain or increase tumor vascularity in preclinical models [12]. While it is difficult to draw broad conclusions from this trial, in the case where extinction of the resistant subpopulation is possible, leveraging the maintained tumor vascularity may provide an avenue for complete tumor eradication. Specifically, if it were known that the resistant population was completely extinct, high-dose cytotoxic treatment could be applied to destroy the residual sensitive cell population.

To our knowledge, this work represents the first direct analytical comparison across dosing schemes between intermittent adaptive therapy, continuous adaptive therapy, and continuous fixed-dose therapy. We additionally show both practical and mathematical advantages of continuous fixed-dose treatment in regions where resistant population extinction is achievable. We expect that this analytical framework will have applications across other adaptive therapy models as well as wider applications in ecology where comparison between discrete control systems and continuous control systems is desirable.

## Methods

Further discussion of the mathematical model and associated proofs can be found in the supplemental material. Simulations were performed using Mathematica [24].

## Acknowledgements

This work was supported by NSF MRSEC Grant (DMR-230934), Simons Foundation Math + X Grant (234606), NIH individual fellowship F30 (5F30CA213737-02), and the Mayo Foundation for Research and Education.

## References

[1] Robert A Gatenby et al. “Adaptive therapy”. In: Cancer research 69.11 (2009), pp. 4894–4903.

[2] Jingsong Zhang et al. “Integrating evolutionary dynamics into treatment of metastatic castrate-resistant prostate cancer”. In: Nature communications 8.1 (2017), pp. 1–9.

[3] Nadia Howlader et al. “The effect of advances in lung-cancer treatment on population mortality”. In: New England Journal of Medicine 383.7 (2020), pp. 640–649.

[4] Angela B Mariotto et al. “Estimation of the number of women living with metastatic breast cancer in the United States”. In: Cancer Epidemiology, Biomarkers & Prevention 26.6 (2017), pp. 809–815.

[5] Ian Smith. “Goals of treatment for patients with metastatic breast cancer”. In: Seminars in oncology. Vol. 33. NElsevier. 2006, pp. 2–5.

[6] Skipper HE. “Implications of biochemical, cytokinetics, pharmacologic, and toxicologic relationships in the design of optimal therapeutic schedules”. In: Cancer Chemother Rep 54 (1970), pp. 431–450.

[7] Douglas Hanahan, Gabriele Bergers, Emily Bergsland, et al. “Less is more, regularly: metronomic dosing of cytotoxic drugs can target tumor angiogenesis in mice”. In: The Journal of clinical investigation 105.8 (2000), pp. 1045–1047.

[8] Irina Kareva, David J Waxman, and Giannoula Lakka Klement. “Metronomic chemotherapy: an attractive alternative to maximum tolerated dose therapy that can activate anti-tumor immunity and minimize therapeutic resistance”. In: Cancer letters 358.2 (2015), pp. 100–106.

[9] Nicolas André, Manon Carré, and Eddy Pasquier. “Metronomics: towards personalized chemotherapy?” In: Nature reviews Clinical oncology 11.7 (2014), pp. 413–431.

[10] Luc G T Morris et al. “Pan-cancer analysis of intratumor heterogeneity as a prognostic determinant of survival.” eng. In: Oncotarget 7.9 (2016), pp. 10051–10063. issn: 1949-2553 (Electronic); 1949-2553 (Linking). doi: 10.18632/oncotarget.7067.

[11] Jingsong Zhang et al. “Evolution-based mathematical models significantly prolong response to abiraterone in metastatic castrate-resistant prostate cancer and identify strategies to further improve outcomes”. In: eLife 11 (2022). Ed. by George H Perry, e76284. issn: 2050-084X. doi: 10.7554/eLife.76284. url: 10.7554/eLife.76284.

[12] Pedro M Enriquez-Navas et al. “Exploiting evolutionary principles to prolong tumor control in preclinical models of breast cancer”. In: Science translational medicine 8.327 (2016), 327ra24–327ra24.

[13] Inna Smalley et al. “Leveraging transcriptional dynamics to improve BRAF inhibitor responses in melanoma”. In: EBioMedicine 48 (2019), pp. 178–190.

[14] Jeffrey West et al. “A survey of open questions in adaptive therapy: Bridging mathematics and clinical translation”. In: Elife 12 (2023), e84263.

[15] Nathan Moore, JeanMarie Houghton, and Stephen Lyle. “Slow-cycling therapy-resistant cancer cells”. In: Stem cells and development 21.10 (2012), pp. 1822–1830.

[16] H J Broxterman et al. “Induction by verapamil of a rapid increase in ATP consumption in multidrug-resistant tumor cells.” eng. In: FASEB J 2.7 (1988), pp. 2278–2282. issn: 0892-6638 (Print); 0892-6638 (Linking). doi:10.1096/fasebj.2.7.3350243.

[17] Maximilian AR Strobl et al. “Turnover modulates the need for a cost of resistance in adaptive therapy”. In: Cancer research 81.4 (2021), pp. 1135–1147.

[18] Jill A Gallaher et al. “Spatial Heterogeneity and Evolutionary Dynamics Modulate Time to Recurrence in Continuous and Adaptive Cancer Therapies.” eng. In: Cancer Res 78.8 (2018), pp. 2127–2139. issn: 1538-7445 (Electronic); 0008-5472 (Print); 0008-5472 (Linking). doi:10.1158/0008-5472.CAN-17-2649.

[19] Yannick Viossat and Robert Noble. “A theoretical analysis of tumour containment.” eng. In: Nat Ecol Evol 5.6 (2021), pp. 826–835. issn: 2397-334X (Electronic); 2397-334X (Linking). doi: 10.1038/s41559-021-01428-w.

[20] L. M. Sonneborn and F. S. Van Vleck. “The Bang-Bang Principle for Linear Control Systems”. In: Journal of The Society for Industrial and Applied Mathematics, Series A: Control 2 (1964), pp. 151–159. url: https://api.semanticscholar.org/CorpusID:123110370.

[21] Alfred J. Lotka. “ELEMENTS OF PHYSICAL BIOLOGY”. In: Science Progress in the Twentieth Century (1919-1933) 21.82 (1926), pp. 341–343. issn: 20594941. url: http://www.jstor.org/stable/43430362 (visited on 08/17/2023).

[22] Jessica Cunningham et al. “Optimal control to reach eco-evolutionary stability in metastatic castrate-resistant prostate cancer”. In: Plos one 15.12 (2020), e0243386.

[23] Masud MA, Jae-Young Kim, and Eunjung Kim. “Containing Cancer with Personalized Minimum Effective Dose”. In: BioRxiv (2022), pp. 2022–03.

[24] Wolfram Research, Inc. Mathematica, Version 14.0. Champaign, IL, 2024. url: https://www.wolfram.com/mathematica.

