## Supplementary Information for "A Mathematical Framework for Comparison of Intermittent versus Continuous Adaptive Chemotherapy Dosing in Cancer"

The supplementary section is meant to provide the mathematical proofs and framework related to the results stated in the main text of this paper. It is divided into the following sections:

2.5 Intermittent Adaptive Therapy converges to Continuous Adaptive Therapy as $\delta \to 0$ 33

### 1 Mathematical Framework

For the ease of the reader, the underlying model and assumptions stated in the main text are described again below.

#### 1.1 Underlying Model

Consider two cell populations. Let the total cell population that is **sensitive** to drug be given by $x$ and the cell population that is **resistant** to drug be given by $y$. Let the **dose** of drug be given by $r_d$. We define $K_A(t)$ to be a function of time $t$ so that $K_A(t) = 0$ when the drug is off and

$K_A(t) = 1$ when the drug is applied. The system of differential equations governing the behavior of $x$ and $y$ can now be written as:

$$\frac{dx}{dt} = f(x,y) + K_A(t)h(x,r_d) \tag{4}$$

$$\frac{dy}{dt} = g(x,y) \tag{5}$$

The effect of the drug on the sensitive population, $x$, is given by $h(x, r_d)$. The function $K_A(t)$ depends on the mode of the therapy, and will be specified below.

In order to model cell-cell competition, we choose $f(x, y)$ and $g(x, y)$ of the form

$$f(x,y) = r_x x(t)(1 - x(t) - \alpha y(t)) \tag{6}$$

$$g(x,y) = r_y y(t)(1 - y(t) - \beta x(t)) \tag{7}$$

where $\alpha, \beta > 0$. A conceptual diagram of this model is shown in Figure 7. Implicit in this set of equations are the following biological assumptions:

- Logistic Growth: The observed growth rates of both cell populations declines as the system reaches carrying capacity.
- Competition: The observed growth rate of each cell population is negatively regulated by the total volume of cells of the same species with a term of 1 and the total volume of cells of the other species with parameter $\alpha$ or $\beta$. For example, if $\alpha > 1$, then species $x$ is more strongly inhibited by the total amount of species y than by the its own species.
- Carrying Capacity: It is assumed that the carrying capacities for $x$ and $y$ (when their respective competitors are absent) are both equal (based on space, resources, etc). The carrying capacity is normalized to 1.

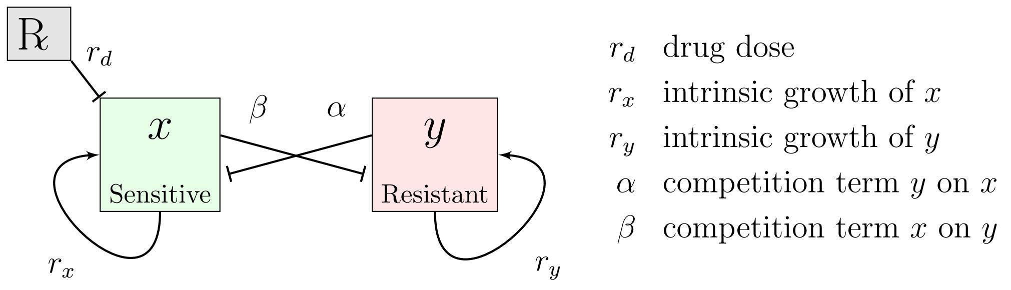

Figure 7: **Conceptual depiction of nonlinear mathematical model.** The sensitive cell population, $x$, and the resistant population $y$ compete with each other and themselves. The drug, ℞, only affects the sensitive cell population, $x$, and is given at some dose $r_d$.

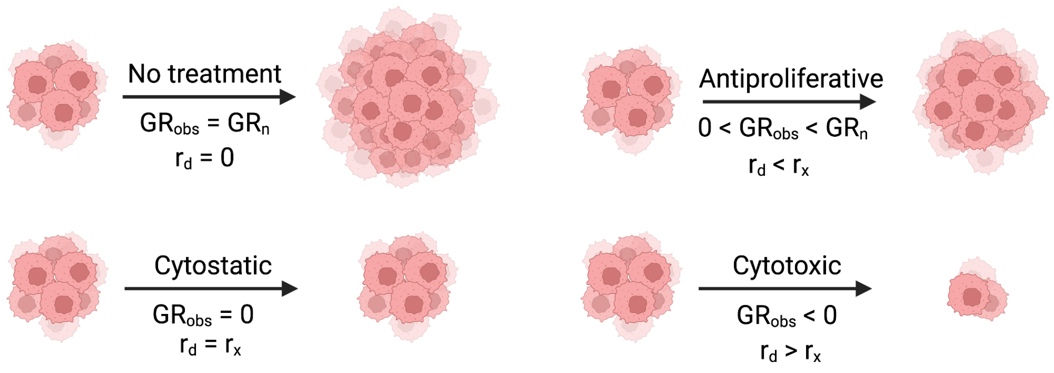

Figure 8: **Pharmacologic Definitions of Drug Action.** $GR_{obs}$ is the observed growth rate of exponentially growing cells under drug treatment and $GR_n$ is the natural exponential growth rate of cells. Created with BioRender.com

#### 1.2 Model Behavior in the Absence of Drug

In the absence of drug ($K_A = 0$), the behavior of this system is dependent on the growth and competition parameters, $r_x, r_y, \alpha, \beta > 0$. The nullclines are calculated below:

$$y = \frac{1-x}{\alpha} \qquad x\text{-nullcline} \tag{8}$$

$$y = 1 - \beta x \qquad y\text{-nullcline} \tag{9}$$

These nullclines are plotted in orange in phase space in Main Figure 2. Note that the intrinsic growth rates $(r_x, r_y)$ do not affect the equilibria or nullclines.

#### 1.3 Model of Drug Action

In order to capture a range of drug effects and retain analytical tractability, a simple model of drug action is considered that allows variable dose control on the sensitive species, $x$. The equation for $h(x, r_d)$ is shown below.

$$h(x, r_d) = -r_d x \tag{10}$$

Consider the action of $h(x, r_d)$ on the single species, $x$, in the absence of resistant species, $y$. Then differential equation for the single species is shown below:

$$\frac{dx}{dt} = r_x x(1-x) - r_d x \tag{11}$$

From a pharmacologic perspective, *in vitro* drug action is measured by comparing relative growth of exponentially dividing cells under varying concentrations of drug, $r_d$. In this model, exponential growth occurs when $x \ll 1$. Thus, a **cytostatic drug** can be approximated as $r_d \approx r_x$, a **cytotoxic drug** as $r_d > r_x$ and an **antiproliferative drug** as $r_d < r_x$ (See Figure 8).

Because these drug action definitions are measured in exponentially growing cells, it is not obvious how to apply these terms appropriately in a logistic system. We note that an antiproliferative agent changes the $x$ equilibrium to

$$x_{eq} = \frac{r_x - r_d}{r_x} \tag{12}$$

Essentially, $x_{eq}$ becomes the new carrying capacity for $x$.

This yields an important observation on drug action near carrying capacity in cellular systems - namely that as a cell population approaches carrying capacity, an antiproliferative agent may appear cytostatic or even cytotoxic (See Figure 5A-C.)

#### 1.4 Modeling Intermittent Adaptive Therapy

Let us now state the model for intermittent adaptive therapy. Since the drug is only effective on the sensitive species, $x$, we aim to use this drug in intervals to preserve the competitive inhibition of $x$ on $y$. A cytotoxic drug ($r_d > r_x$) is given when the tumor volume $x(t) + y(t)$ reaches a certain threshold, $A > 0$, and is then discontinued when the tumor volume hits a defined lower threshold $A - \delta > 0$ where $\delta > 0$. The tumor is then allowed to regrow until it reaches volume $A$ at which point the drug is reapplied. Adaptive therapy is thus modeled as a bang-bang controller switching abruptly between drug off and drug on states depending on the tumor volume $x(t) + y(t)$. A key modeling assumption is that one indeed has complete real-time knowledge of total tumor volume. The tumor volume can be measured *in vitro* directly or *in vivo* using imaging or biomarkers [2, 17, 1].

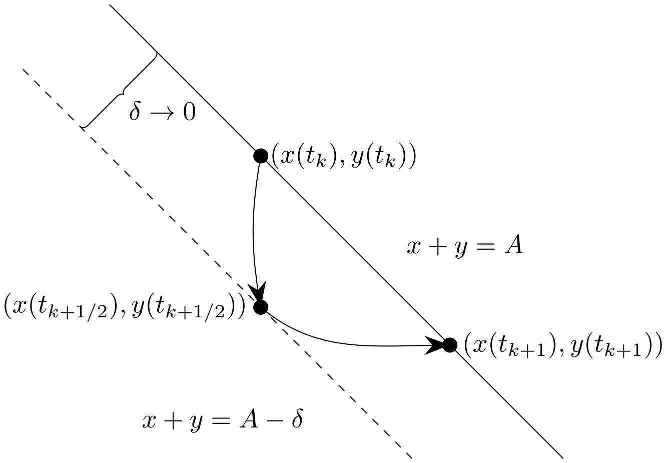

Figure 9: Illustration of flow in intermittent adaptive therapy. Solid line: $x + y = A$, broken line: $x + y = A - \delta$.

Formally, we may describe intermittent therapy as follows (see Figure 9). We start at time $t = 0$. Suppose $x(0) + y(0) > A$. Then, we set $K_A(t) = 1$ in (4) for $0 \le t < t_0$, where $t_0$ will be specified below. If $x(0) + y(0) < A$, we set $K_A(t) = 0$ for $0 \le t < t_0$. Let $t = t_0$ be the first time at which:

$$x(t_0) + y(t_0) = A. \tag{13}$$

If there is no such $t_0$, we set $t_0 = \infty$. If $x(0) + y(0) = A$, set $t_0 = 0$. At $t = t_0$, we turn on therapy, so that $K_A(t) = 1$ for $t_0 \le t < t_{1/2}$, where we specify $t_{1/2}$ as follows. Let $t = t_{1/2}$ be the first time for $t > t_0$ at which

$$x(t_{1/2}) + y(t_{1/2}) = A - \delta. \tag{14}$$

If no such $t_{1/2}$ exists, we set $t_{1/2} = \infty$. At $t = t_1$, we turn off therapy so that $K_A(t) = 0$ for $t_{1/2} \le t < t_1$, where we specify $t_1$ as follows. Let $t = t_1$ be the first time $t > t_{1/2}$ at which:

$$x(t_1) + y(t_1) = A. \tag{15}$$

If such a $t_1$ does not exist, we set $t_1 = \infty$. We then switch on therapy again so that $K_A(t) = 1$. This process is repeated. We thus have a sequence of switching times $t_k, t_{k+1/2}, k \in \mathbb{Z}, k \ge 0$ so that:

$$K_A(t) = \begin{cases} 1 & \text{if } t_k \le t < t_{k+1/2}, \\ 0 & \text{if } t_{k+1/2} \le t < t_{k+1}. \end{cases} \tag{16}$$

For $t_{k+1/2}$ to exist, the tumor volume $x(t) + y(t)$ must drop to $A - \delta$ after the drug is turned on at $t = t_k$. If such a $t_{k+1/2}$ does not exist, we set $t_{k+1/2} = \infty$, and all subsequent $t_\ell, \ell > k + 1/2$ are not defined. Likewise, for $t_{k+1}$ to exist, tumor volume must increase back to $A$ after which the drug is turned off at $t = t_{k+1/2}$. If such a $t_{k+1}$ does not exist, we set $t_{k+1} = \infty$, and all subsequent $t_\ell, \ell > k + 1$ are not defined.

We assume that $A < 1$ (the carrying capacity). The reason for this assumption is because if $A \ge 1$ in the phase spaces shown in Figure 2, in most cases the optimal therapy would be no therapy.

### 2 Analysis of Intermittent Adaptive Therapy

#### 2.1 Dynamics of Intermittent Adaptive Therapy

Using the model described in Section 1, the phase space of intermittent adaptive therapy is explored in two cases. The first case considered is the case of a fitness cost associated with resistance where $x$ is a stronger competitor to $y$ and $\alpha < 1$ and $\beta > 1$. In the second case, the application of adaptive therapy when the resistant species is the stronger competitor ($\alpha > 1, \beta < 1$) is explored.

##### 2.1.1 Case 1: $\alpha < 1, \beta > 1$

The case where the sensitive species is the stronger competitor (fitness cost) is described in detail in the main text (Section ). In brief, the control boundaries ($x + y = A$ and $x + y = A - \delta$) are two parallel lines in $x, y$ phase space that intersect with the $y$-nullcline which is fixed and independent of the presence or absence of drug. The $x$-nullcline is to the right of the control lines under no drug action ($x$ is always increasing) and to the left of the control lines under cytotoxic drug action ($x$ is always decreasing). In Figure 3, we can see that there are two generic scenarios: resistant population extinction (Figure 3D) and sensitive population extinction (Figure 3B). In the case of resistant population extinction, the sequence of switching times $t_k, t_{k+1/2}$ defined in (13), (14), (15) and (16) are defined for all $k \in \mathbb{Z}, k \ge 0$. However, for the case of sensitive population extinction, the switching times are not defined beyond a certain point. In Figure 3B, the last switching time is $t_1$ ($t_{3/2} = \infty$). The two generic behaviors are separated by an unstable limit cycle shown in Figure 3C.

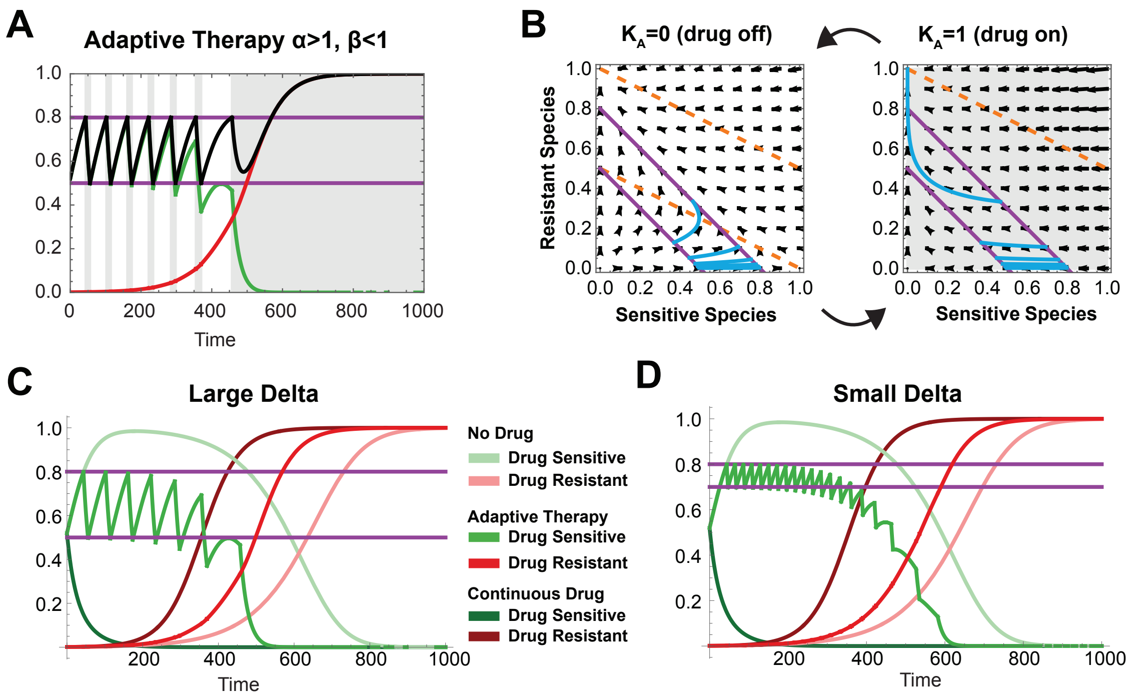

Figure 10: **Adaptive therapy when $\alpha > 1, \beta < 1$.** (**A**) Time course of resistant outgrowth during adaptive therapy with phase space plotted in (**B**). (**C,D**) Comparison of adaptive therapy, no therapy, and fixed dose continuous cytotoxic therapy with wide control (large delta) and tight control (small delta).

##### 2.1.2 Case 2: $\alpha > 1, \beta < 1$

When $\alpha > 1$ and $\beta < 1$, the resistant species always out competes the sensitive species and resistant species eradication is not achievable under adaptive therapy. A simulation is shown in Figure 10A,B. Note that the switching times are only well-defined up to a certain point.

While there is no strategy in which the tumor volume is kept indefinitely below $A$, adaptive therapy lengthens the time until the tumor volume progresses above $A$ when compared with continuous dosing of the same dose of drug or no therapy (Figure 10C,D). This effect is due to competitive inhibition of the sensitive species, $x$, on the growth of $y$ and has been described previously [17]. The magnitude of the difference between the time to tumor progression of adaptive therapy versus continuous high dose therapy is dependent on $\delta$. Since $y$ is monotonically increasing and $\frac{dy}{dt}$ is negatively regulated by $\beta x$, the higher the value of $x$, the slower the growth of $y$. Similarly, the slowest time to resistant outgrowth of the tumor at volume greater than $A$ occurs under no treatment.

Overall adaptive therapy provides a useful method for delaying resistant population outgrowth while keeping the total tumor volume under a given threshold, $A$.

#### 2.2 Limiting Dynamics on the $x + y = A$ Line and Continuous Adaptive Therapy

It is, in general, difficult to mathematically analyze the dynamics of intermittent adaptive therapy. However, when $\delta$ is small, we can obtain a complete understanding of the dynamics. This can be seen from the following observation. As $\delta$ becomes smaller, the dynamics of intermittent adaptive therapy is increasingly confined to a narrow window around the line $x + y = A$. This is also graphically seen in Figure 3 and Figure 10. If we turn our attention to the defining equations (4) and (5), we notice that (5) does not change whether or not there is adaptive therapy. In the limit at $\delta \to 0$, then, our expectation is that the limiting system may be equivalent to:

$$\frac{dy}{dt} = g(x, y), \text{ and } x + y = A. \tag{17}$$

That is to say, in the limit as $\delta \to 0$, the effect of (4) is merely to enforce the condition $x + y = A$. Now, consider the equation:

$$\frac{dz}{dt} = g(A - z, z), \ z(0) = z_0. \tag{18}$$

The expectation is that the solution $z(t)$ will give a good approximation to $y(t)$. This expectation is indeed correct, under certain conditions. Our first result in this direction is the following.

**Proposition 2.1.** *For fixed time, $T > 0$, let $\{x(t), y(t)\}$ be solutions to the system of differential equations for some initial conditions $(x(0), y(0)) = (x_0, y_0), x_0 + y_0 = A$ such that $A - \delta \leq x(t) + y(t) \leq A$ for $0 \leq t \leq T$. Let $z(t)$ be the solution to (18) with $z(0) = z_0 = y_0$. Then, there is a constant $C$ that only depends only on $T$ such that*

$$|y(t) - z(t)| \leq C\delta \text{ for } 0 \leq t \leq T. \tag{19}$$

*Proof.* We begin by examining the differential equation for $w = y - z$:

$$\begin{aligned}\frac{dw}{dt} &= \frac{d(y-z)}{dt} = \frac{dy}{dt} - \frac{dz}{dt} \\ &= g(x, y) - g(A - z, z) \\ &= g(x, y) - g(A - y, y) + g(A - y, y) - g(A - z, z)\end{aligned} \tag{20}$$

Equation 20 can now be split into two parts $R(t)$ and $c(t)w$ for which bounds will be evaluated separately.

$$R(t) = g(x, y) - g(A - y, y) \tag{21}$$

$$c(t)w = g(A - y, y) - g(A - z, z) \tag{22}$$

Examining Equation 21, we note that

$$R(t) = \int_{A-y}^{x} \frac{\partial g}{\partial x}(s, y)ds.$$

We can now bound $|R(t)|$ by noting that $\frac{\partial g}{\partial x} = -r_y y\beta$, $y \leq A$, and $|x - (A - y)| \leq \delta$ to yield:

$$|R(t)| \leq \delta r_y A\beta \tag{23}$$

Next, we examine Equation 22. Let $h(y) = g(A-y, y)$, then

$$g(A-y,y) - g(A-z,z) = h(y) - h(z) = (y-z)c(t)$$

where

$$c(t) = \int_0^1 h'(sy + (1-s)z)ds \tag{24}$$

We can determine an upper bound of $|c(t)|$ by directly evaluating

$$\frac{dh}{dy} = \frac{d(g(A-y,y))}{dy} = r_y(1-A\beta) + 2r_y y(\beta - 1)$$

Since $y \le A$ and all other terms are bounded, we let $L$ be defined such that $|r_y(1-A\beta) + 2r_y y(\beta-1)| \le L$, then

$$|c(t)| \le L \tag{25}$$

We can now write Equation 20 as

$$\frac{dw}{dt} - c(t)w = R(t). \tag{26}$$

The equation for $w(t)$ can be solved exactly using a general linear equation. We first multiply both sides by $I(t) = e^{-\int_0^t c(s)ds}$ to yield

$$e^{-\int_0^t c(s)ds}\left(\frac{dw}{dt} - c(t)w\right) = R(t)e^{-\int_0^t c(s)ds}$$

Appealing to the chain rule, we note that this can be re-written as

$$\frac{d}{dt}\left(w(t)e^{-\int_0^t c(s)ds}\right) = R(t)e^{-\int_0^t c(s)ds}.$$

Integrating both sides yields:

$$w(t)e^{-\int_0^t c(s)ds} - w(0) = \int_0^t R(\sigma)e^{-\int_0^\sigma c(s)ds}d\sigma$$

We note that $w(0) = y(0) - z(0) = 0$, so we can solve for $w(t)$

$$w(t) = e^{\int_0^t c(s)ds}\int_0^t R(\sigma)e^{-\int_0^\sigma c(s)ds}d\sigma$$

For any fixed $t < T$, $e^{\int_0^t c(s)ds}$ is constant, so

$$\begin{aligned} w(t) &= \int_0^t R(\sigma)e^{\int_0^t c(s)ds - \int_0^\sigma c(s)ds}d\sigma \\ &= \int_0^t R(\sigma)e^{\int_\sigma^t c(s)ds}d\sigma \end{aligned} \tag{27}$$

We have already established a bound on $|R(\sigma)|$ in equation 23 and a bound on $c(s)$ in 25,

$$|w(t)| \leq \delta r_y A\beta \int_0^t e^{L(t-\sigma)} d\sigma = \frac{\delta r_y A\beta(e^{Lt}-1)}{L} \leq C(T)\delta, \tag{28}$$

$$C(T) = \frac{\delta r_y A\beta(e^{LT}-1)}{L}. \tag{29}$$

This establishes (19). □

The above result suggests that, when $\delta$ is small, the dynamics of intermittent adaptive therapy can be understood by studying (18), which is considerably easier to understand. For example, it is a trivial matter to determine the stable and unstable fixed points of (18). However, the above result rests on the crucial assumption that the value of $x + y$ lies between $A - \delta$ and $A$ over some time period $T$. We do not know how $T$ may depend on $\delta$; indeed, $T$, the time period over which the above assumption is valid, may vanish as $\delta \to 0$. To show that the properties of the dynamical system (18) do indeed correspond to the dynamics of intermittent adaptive therapy as $\delta \to 0$, we need to look more closely at the dynamics of intermittent adaptive therapy.

Before we proceed, let us point out that the limiting dynamics described by (17) (or equivalently, (18)) may be interpreted as that of continuous adaptive therapy. Note that, as long as $x + y = A$, we have:

$$\frac{dx}{dt} + \frac{dy}{dt} = 0.$$

Since $y$ satisfies (5), we thus have:

$$\frac{dx}{dt} = -g(x, y).$$

Now, let us consider the possibility of achieving this dynamics by modulating the drug dose (see (4) and (10)).

$$\frac{dx}{dt} = -g(x,y) = f(x,y) - \widehat{R}_c(x,y)x, \quad \widehat{R}_c(x,y) = \frac{f(x,y)+g(x,y)}{x}. \tag{30}$$

What we see is that the limiting dynamics on the line $x+y = A$ can be achieved by letting the drug dose equal to $\widehat{R}_c(x, y)$ defined above. The drug level $\widehat{R}_c(x, y)$ must be positive, so we must have $f(x, y)+g(x, y) > 0$ for this to make sense. With this in mind, we introduce the following idealized version of adaptive therapy, which we term *continuous adaptive therapy.* In continuous adaptive therapy, we perform the following. If the initial condition $x(0), y(0)$ satisfies $x(0) + y(0) > A$, we turn the drug on so that $K_A(t) = 1$ for $0 < t < t_0$. If $x(0) + y(0) < A$, we set $K_A(t) = 0$ for $0 < t < t_0$. At $t = t_0$, we hit $x(t_0) + y(t_0) = A$. For $t \geq t_0$, we turn our drug on at the level of $\widehat{R}_c(x, y)$ so long as $\widehat{R}_c(x, y)$ is positive. We will not define continuous adaptive therapy in regions where $\widehat{R}_c(x, y) \leq 0$ (or equivalently, $f(x, y) + g(x, y) \leq 0$).

What Proposition 2.1 suggests is that as $\delta \to 0$, intermittent adaptive therapy approaches continuous adaptive therapy.

#### 2.3 Intermittent Adaptive Therapy and the Discrete Map $\mathcal{F}_\delta$

As seen above, intermittent adaptive therapy bounds the tumor size to be between $A - \delta$ and $A$, assuming that the tumor size decreases sufficiently in the presence of drug and increases sufficiently in the absence of drug. To better understand this behavior, we introduce some definitions. First, we introduce some maps (see Figure 11). Take any point $(A-y,y)$. Let $z=\Phi_\delta(y)$ be the map that takes $(A-y,y)$ to $(A-\delta-z,z)$ under the flow when the drug is on. More precisely, let $(\widehat{x}(t),\widehat{y}(t))$ be the solution to (4) and (5) with $K_A(t)=1$ and initial data $(\widehat{x}(0),\widehat{y}(0))=(A-y,y)$. Suppose there is a $\widehat{t}_*$ such that

$$A-\delta<\widehat{x}(t)+\widehat{y}(t)<A \text{ for } 0<t<\widehat{t}_* \text{ and } \widehat{x}(\widehat{t}_*)+\widehat{y}(\widehat{t}_*)=A-\delta. \tag{31}$$

Then, let $\Phi_\delta(y)\equiv\widehat{y}(\widehat{t}_*)$ and $\mathcal{T}_\delta^\Phi(y)\equiv\widehat{t}_*$. Likewise, take any point $(A-\delta-y,y)$. Let $z=\Psi_\delta(y)$ be that map that takes $(A-y-\delta,y)$ to $(A-z,z)$ under the flow when the drug is off. If we let $(\widetilde{x}(t),\widetilde{y}(t))$ be the solution to (4) and (5) with $K_A(t)=0$ and initial data $(\widetilde{x}(0),\widetilde{y}(0))=(A-y-\delta,y)$. Suppose there is a $\widetilde{t}_*$ such that

$$A-\delta<\widetilde{x}(t)+\widetilde{y}(t)<A \text{ for } 0<t<\widetilde{t}_* \text{ and } \widetilde{x}(t_*)+\widetilde{y}(\widetilde{t}_*)=A.$$

Then, let $\Psi_\delta(y)\equiv\widetilde{y}(\widetilde{t}_*)$ and $\mathcal{T}_\delta^\Psi(y)\equiv\widetilde{t}_*$. Let us define the composite maps $\mathcal{F}_\delta(y)=\Psi_\delta(\Phi_\delta(y))$ and $\mathcal{T}_\delta(y)=\mathcal{T}_\delta^\Phi(y)+\mathcal{T}_\delta^\Psi(\Phi_\delta(y))$. None of the above maps are necessarily well-defined for all values of $y$ or $\delta$. However, if these maps are well-defined, we can relate them to intermittent adaptive therapy as follows. Let $x=x_\delta(t)$ and $y=y_\delta(t)$ be the solutions intermittent adaptive therapy dynamics with the high and low tumor volume thresholds equal to $A$ and $A-\delta$. Then,

$$y_\delta(t_{k+1})=\mathcal{F}_\delta(y_\delta(t_k)),\ t_{k+1}-t_k=\mathcal{T}_\delta(y_\delta(t_k)). \tag{32}$$

Thus, so long as $\mathcal{F}_\delta$ is well-defined at $y(t_k)$ (in which case $\mathcal{T}_\delta$ is also well-defined), studying intermittent adaptive therapy reduces to studying these discrete maps.

**Definition 2.1** (Weak Control Region)**.** *A bounded open interval $a<y<b$ for which $\mathcal{F}_\delta(y)$ is well-defined is a weak control region.*

**Definition 2.2** (Strict Control Region)**.** *Let $\mathcal{I}$ be a weak control interval. If $y\in\mathcal{I}$ implies $\mathcal{F}_\delta(y)\in\mathcal{I}$, then $\mathcal{I}$ is a strict control region.*

In other words, $\mathcal{F}_\delta$ is well-defined on a weak control region, and any region that is invariant under the action of $\mathcal{F}_\delta$ is a strict control region. In a strict control region, the switching times $t_k$ will be defined for all $k$. It is in general difficult to analytically identify weak or strict control regions. It is, however, possible to find these regions when $\delta$ is small. For this, we introduce the notion of control region. We consider the vector field defined by the differential equation (4) and (5) when $K_A=1$ or $K_A=0$, and see if the vector field on the line $x+y=A$ is pointing in the direction of increase or decrease of tumor size.

**Definition 2.3** (Control Region)**.** *Consider the vector fields $(f+h,g)$ and $(f,g)$ characterizing the differential equations* (4) *and* (5)*. The values of $y$ for which the following inequalities hold is the control region:*

$$-p_A(y)\equiv f(A-y,y)+h(A-y,r_d)+g(A-y,y)<0, \tag{33}$$

$$q_A(y)\equiv f(A-y,y)+g(A-y,y)>0. \tag{34}$$

Control regions can be easily identified by examining the above inequalities. Let the *area of controllability* $\mathcal{C}$ be the subset of the $(x,y)\in\mathbb{R}^2$ satisfying:

$$f(x,y)+h(x,r_d)+g(x,y)<0 \text{ and } f(x,y)+g(x,y)>0. \tag{35}$$

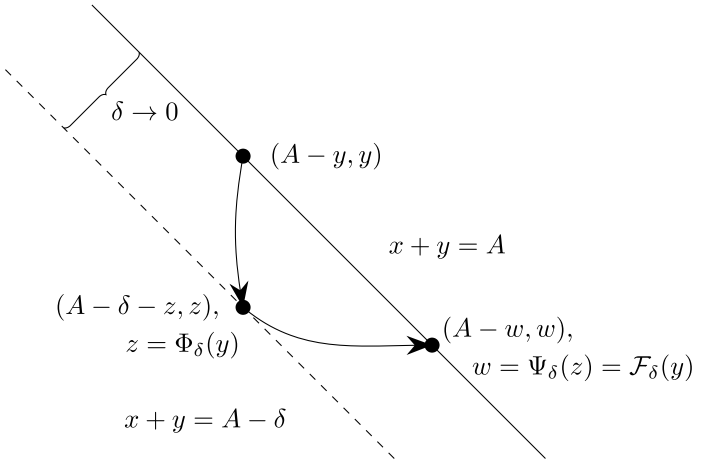

Figure 11: Illustration of the maps $\Phi_\delta, \Psi_\delta$ and $\mathcal{F}_\delta$. Solid line: $x+y=A$, broken line: $x+y=A-\delta$.

This region is shown graphically in Figure 12. The portion of the line $x+y=A$ that is contained in $\mathcal{C}$ corresponds to the control region. We also note that, as long as $x>0$, the first condition above is equivalent to the statement that:

$$r_d > \widehat{R}_c(x,y) = \frac{f(x,y)+g(x,y)}{x}, \tag{36}$$

where $R_c$ is the drug level for continuous adaptive therapy defined in (30). The relationship between the control region and weak control region is given by the following result.

**Proposition 2.2.** *Consider any open interval $\mathcal{I}$ such that its closure is contained in the control region. Then, there is a $\delta_*>0$ such that for all $\delta<\delta_*$, $\mathcal{I}$ is a weak control region.*

*Sketch of Proof.* The above result is consequence of smooth dependence of solutions to differential equations on initial data, together with the implicit function theorem. Let $(x(t),y(t))$ be the solution to the differential equation (4) and (5) with initial data $(x(0),y(0))=(x_0,y_0)$ and $K_A=1$. Define the flow map:

$$\phi_t(x_0,y_0) = \begin{pmatrix} \phi_x(x_0,y_0,t) \\ \phi_y(x_0,y_0,t) \end{pmatrix} = \begin{pmatrix} x(t) \\ y(t) \end{pmatrix}$$

It is well-known that both $\phi_x$ and $\phi_y$ are smooth functions of $(x_0,y_0,t)$ in any open set on which $\phi_x$ and $\phi_y$ are defined. Let:

$$G(y,t,\delta) = \phi_x(A-y,y,t) + \phi_y(A-y,y,t) - (A-\delta). \tag{37}$$

Note that:

$$G(y,0,0)=0.$$

Suppose $y\in\mathcal{I}$. Then,

$$\frac{\partial G}{\partial t}(y,0,0) = -p_A(y) = f(A-y,y) + h(A-y,r_d) + g(A-y,y) < 0$$

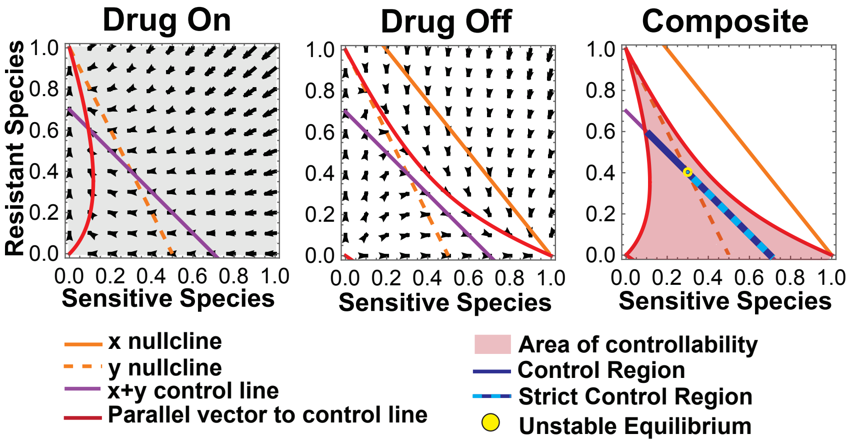

Figure 12: **Continuous control, strict control region, and control region for $\alpha < 1$ and $\beta > 1$.**

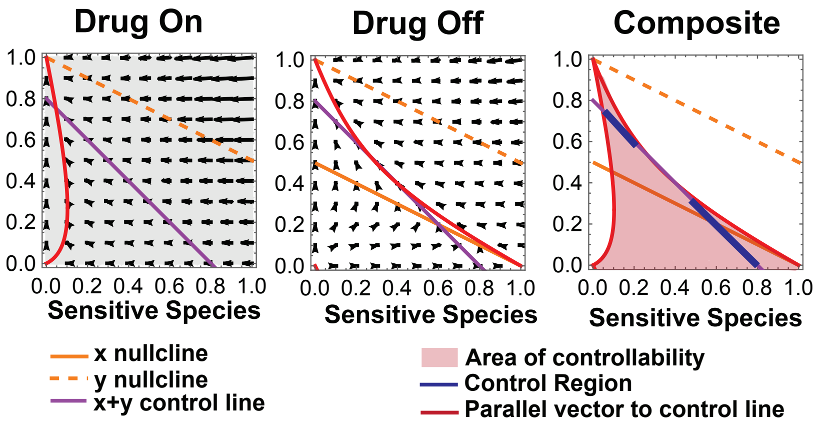

Figure 13: **Continuous control, strict control region, and control region for $\alpha < 1$ and $\beta > 1$.**

where we used the definition of control region. We may thus use the implicit function theorem to solve for $t = t_\phi(\delta, y)$ in terms of $y$ and $\delta$ for $|\delta| < \delta_*, \delta_* > 0$. We can take $\delta_*$ to be independent of $y$ by the assumption that the closure of $\mathcal{I}$ is a compact set contained within the control region. We may thus let $\Phi_\delta(y) = \phi_y(A - y, y, t_\phi(y, \delta))$. It is clear that $\phi_t(A - y, y)$ satisfies the inequality in (31) when $0 < t < t_\phi(y, \delta)$. Take an open interval $\mathcal{J}$ that contains the closure $\mathcal{I}$, such that the closure of $\mathcal{J}$ is still contained in the control region. This is always possible since the control region is an open set. In much the same way as above, $\Psi_\delta(y)$ can be defined on $\mathcal{J}$ for $|\delta| < \delta_*, \delta_* > 0$, by possibly making $\delta_*$ even smaller. By taking $\delta_*$ smaller still if necessary, we can make sure that $y \in \mathcal{I}$ implies $\Phi_\delta(y) \in \mathcal{J}$. This implies that $\mathcal{F}_\delta(y) = \Psi_\delta(\Phi_\delta(y))$ is defined on $\mathcal{I}$ for $|\delta| < \delta_*$. □

Note that the above proof gives a construction of $\mathcal{F}_\delta(y)$ that is defined for $|\delta| < \delta_*$ (not only $\delta > 0$) and $y \in \widehat{\mathcal{I}}$, where $\widehat{\mathcal{I}}$ is some open interval that includes the closure of $\mathcal{I}$. Furthermore, $\mathcal{F}_\delta(y)$ is a smooth function of $(y, \delta)$ with bounded partial derivatives with respect to $\delta$ and $y$ with and $\mathcal{F}_0(y) = y$.

The following is a technical result that will be useful for us later.

**Lemma 2.3.** *Let $\mathcal{I}$ an interval whose closure is contained in the control region. Then,*

$$\begin{aligned} \lim_{\delta \to 0} \frac{1}{\delta} \left(\mathcal{F}_\delta(y) - y\right) &= g(A - y, y) \left(\frac{1}{p_A(y)} + \frac{1}{q_A(y)}\right) \\ \lim_{\delta \to 0} \frac{1}{\delta} \mathcal{T}_\delta(y) &= \frac{1}{p_A(y)} + \frac{1}{q_A(y)} \ \textit{for } y \in \mathcal{I}, \end{aligned} \tag{38}$$

*where $p_A$ and $q_A$ were defined in (33) and (34). Consider the set $\mathcal{K} = \{(\delta, y) | \, |\delta| < \delta_*, y \in \mathcal{I}\}$. There is a $\delta_*$ small enough such that the functions $(\mathcal{F}_\delta(y) - y)/\delta$ are $\mathcal{T}_\delta/\delta$ are smooth on $\mathcal{K}$. Moreover all partial derivatives $\frac{\partial}{\partial \delta}^k \frac{\partial}{\partial y}^l \left((\mathcal{F}_\delta(y) - y)/\delta\right), \frac{\partial}{\partial \delta}^k \frac{\partial}{\partial y}^l (\mathcal{T}_\delta/\delta)$ $k \geq 0, l \geq 0$ are bounded on $\mathcal{K}$.*

*Sketch of proof.* Consider equation (37) and its solution $t_*(y, \delta)$. By the implicit function theorem, we have:

$$\left. \frac{\partial \mathcal{T}_\delta^\Phi}{\partial \delta}(y) \right|_{\delta=0} = \frac{\partial t_*}{\partial \delta}(y, 0) = -\frac{\partial G / \partial \delta}{\partial G / \partial t}(y, 0, 0) = \frac{1}{p_A(y)}. \tag{39}$$

Thus,

$$\left. \frac{\partial}{\partial \delta} \Phi_\delta(y) \right|_{\delta=0} = \left. \frac{\partial}{\partial \delta} \phi_y(A - y, y, t_*(y, \delta)) \right|_{\delta=0} = g(A - y, y) \frac{\partial t_*}{\partial \delta}(y, 0) = \frac{g(A - y, y)}{p_A(y)}. \tag{40}$$

For $\Psi_\delta(y)$, it is easier to consider the inverse map $\Psi_\delta^{-1}(y)$ instead. Applying the same technique to $\Psi_\delta^{-1}$ as above and using the fact that $\Psi_\delta$ is its inverse map, we obtain:

$$\left. \frac{\partial \mathcal{T}_\delta^\Psi}{\partial \delta}(y) \right|_{\delta=0} = \frac{1}{q_A(y)}, \ \left. \frac{\partial}{\partial \delta} \Psi_\delta(y) \right|_{\delta=0} = \frac{g(A - y, y)}{q_A(y)}. \tag{41}$$

Thus,

$$\begin{aligned} \left. \frac{\partial \mathcal{F}_\delta}{\partial \delta}(y) \right|_{\delta=0} &= \left. \frac{\partial \Psi_\delta}{\partial \delta}(\Phi_0(y)) \right|_{\delta=0} + \frac{\partial \Psi_0}{\partial y}(\Phi_0(y)) \left. \frac{\partial \Phi_\delta}{\partial \delta}(y) \right|_{\delta=0} \\ &= g(A - y, y) \left(\frac{1}{p_A(y)} + \frac{1}{q_A(y)}\right). \end{aligned}$$

Since $\mathcal{F}_0(y) = y$, we obtain the desired limit for $\mathcal{F}_\delta$. The limit for $\mathcal{T}_\delta$ can be similarly computed. The statement about smoothness and partial derivatives follows from the smoothness of the flow maps used in Proposition 2.2. □

#### 2.4 Fixed Points and Invariant Regions of $\mathcal{F}_\delta$ when $\delta$ is small

Recall that Proposition 2.1 suggested that the the dynamics of intermittent adaptive therapy can be studied using the dynamics of (18) (continuous adaptive therapy). Suppose $z = z_*$ is a steady state of (18):

$$g(A - z_*, z_*) = 0.$$

Then, $z = z_*$ is a stable/unstable fixed point if $\frac{d}{dz} g(A-z,z)\Big|_{z=z_*}$ is negative/positive. It is expected that, for positive but small values of $\delta$, the map $\mathcal{F}_\delta(y)$ has a fixed point near $y = z_*$ and that it will be a stable/unstable fixed point depending on whether $z = z_*$ is a stable/unstable fixed point of (18). The following theorem states that this is indeed the case. Note that a fixed point of $\mathcal{F}_\delta$ corresponds to a limit cycle for intermittent adaptive therapy.

**Theorem 2.4.** *Suppose $y = y_0$ is in the control region and:*

$$g(A - y_0, y_0) = 0, \ \frac{d}{dy} g(A-y,y)\Big|_{y=y_0} \neq 0.$$

*Then, there is a $\delta_* > 0$ such that for all $|\delta| < \delta_*$, there is a $y_\delta$ with the following properties.*

1. *$\mathcal{F}_\delta(y_\delta) = y_\delta$ where $y_{\delta=0} = y_0$ and $y_\delta$ is a smooth function of $\delta$. There is an $\epsilon_* > 0$ independent of $|\delta| < \delta_*$ such that $y_\delta$ is the only solution to $\mathcal{F}_\delta(y) = y$ satisfying $|y - y_0| < \epsilon_*$.*

2. *If $\frac{d}{dy} g(A-y,y)\Big|_{y=y_0} > 0$, then $y_\delta$ is an unstable fixed point of the discrete dynamical system defined by $\mathcal{F}_\delta$. If $\frac{d}{dy} g(A-y,y)\Big|_{y=y_0} < 0$, then $y_\delta$ is a stable fixed point.*

3. *$y_\delta$ has the following expansion with respect to $\delta$:*

$$y_\delta = y_0 + \frac{\frac{\partial g}{\partial x}(A - y_0, y_0)}{2\left(\frac{\partial g}{\partial y}(A-y_0,y_0) - \frac{\partial g}{\partial x}(A-y_0,y_0)\right)}\delta + \mathcal{O}(\delta^2). \tag{42}$$

*Proof.* Take an interval $\mathcal{I}$ around $y_0$ so that this interval is in the control region. For $y \in \mathcal{I}$, consider the function:

$$Q(y,\delta) = \frac{1}{\delta}\left(\mathcal{F}_\delta(y) - y)\right)$$

where we set $Q(y,0)$ to be the limiting value given in (38). Note that, for $\delta \neq 0$, $\mathcal{F}_\delta(y) = y$ if and only if $Q(y,\delta) = 0$. It is clear that $Q(y,\delta)$ is a smooth function of $y$ and $\delta$ for sufficiently small $\delta$. We now apply the implicit function theorem on $Q(y,\delta)$. By assumption,

$$\begin{aligned} Q(y_0, 0) &= g(A-y_0,y_0)\left(\frac{1}{p_A(y_0)} + \frac{1}{q_A(y_0)}\right) = 0, \\ \frac{\partial Q}{\partial y}(y_0,0) &= \frac{d}{dy} g(A-y,y)\Big|_{y=y_0}\left(\frac{1}{p_A(y_0)} + \frac{1}{q_A(y_0)}\right) \neq 0. \end{aligned} \tag{43}$$

Thus, item 1. holds by the implicit function theorem. Furthermore,

$$\lim_{\delta\to 0}\frac{1}{\delta}\left(\left.\frac{\partial \mathcal{F}_\delta}{\partial y}\right|_{y=y_\delta}-1\right)=\lim_{\delta\to 0}\frac{\partial Q}{\partial y}(y_\delta,\delta)=\frac{\partial Q}{\partial y}(y_0,0).$$

Since $y_0 \in \mathcal{I}$, $p_A(y_0) > 0$ and $q_A(y_0) > 0$. Equation (43) implies that $\frac{\partial Q}{\partial y}(y_0, 0)$ has the same sign as $\left.\frac{d}{dy}g(A-y,y)\right|_{y=y_0}$. Thus, for sufficiently small $\delta$, $\left.\frac{\partial \mathcal{F}_\delta}{\partial y}\right|_{y=y_\delta}$ is greater or less than 1 depending on whether $\left.\frac{d}{dy}g(A-y,y)\right|_{y=y_0}$ is positive or negative. This proves item 2.

To prove (42), we seek an expansion of $\Phi_\delta$ and $\Psi_\delta$ with respect to $\delta$:

$$\Phi_\delta(y) = y + \delta A_1(y) + \delta^2 A_2(y) + \cdots,$$
$$\Psi_\delta(y) = y + \delta B_1(y) + \delta^2 B_2(y) + \cdots.$$

A straightforward computation shows that:

$$\mathcal{F}_\delta(y) = y + \delta(A_1 + B_1) + \delta^2\left(A_2 + B_2 + \frac{dB_1}{dy}A_1\right) + \cdots. \tag{44}$$

Since $y_\delta$ is a smooth function of $\delta$ with $y_{\delta=0}$, $y_\delta$ has the expansion:

$$y_\delta = y_0 + \delta w + \cdots.$$

Plugging this expression into $\mathcal{F}_\delta(y_\delta) = y_\delta$ and using (44), we see that:

$$w = -\frac{A_2(y_0) + B_2(y_0) + \left.\frac{dB_1}{dy}A_1\right|_{y=y_0}}{\left.\frac{d}{dy}(A_1 + B_1)\right|_{y=y_0}}. \tag{45}$$

Note that $A_1$ and $B_1$ are given respectively by (40) and (41). Note furthermore that $A_1 = B_1 = 0$ at $y = y_0$. Therefore, we have only to compute $A_2(y_0)$ and $B_2(y_0)$. Let us compute $A_2(y_0)$. Consider the solution to the differential equation:

$$\frac{dx}{dt} = \widehat{f}(x,y) = f(x,y) + h(x,r_d),\ \frac{dy}{dt} = g(x,y),\ x(0) = x_0 = A - y_0,\ y(0) = y_0.$$

Since $g(x_0, y_0) = 0$, the orbit $(x(t), y(t))$ is locally a graph that can be expressed as $y(t) = q(x(t))$. We may expand:

$$q(x) = y_0 + \frac{dq}{dx}(x_0)(x-x_0)^2 + \frac{1}{2}\frac{d^2q}{dx^2}(x_0)(x-x_0)^2 + \cdots$$

Since $g(x_0, y_0) = 0$, we see that $\frac{dq}{dx}(x_0) = 0$. After some calculation, we see that:

$$\frac{d^2q}{dx^2}(x_0) = \left.\frac{d^2y}{dt^2}\right|_{t=0}\left(\left.\frac{dx}{dt}\right|_{t=0}\right)^{-2} = \frac{\frac{\partial g}{\partial x}(x_0,y_0)}{\widehat{f}(x_0,y_0)}. \tag{46}$$

We thus have:

$$q(x) = y_0 + \frac{\frac{\partial g}{\partial x}(x_0,y_0)}{2\widehat{f}(x_0,y_0)}(x-x_0)^2 + \cdots.$$

The value of $\Phi_\delta(y_0)$ is the intersection point of $y = q(x)$ and the line $x + y = A - \delta$. Thus, given the value $x_*$ that satisfies $x_* + q(x_*) = A - \delta$, $\Phi_\delta(y_0) = q(x_*)$. From this, it is easily seen that:

$$\Phi_\delta(y_0) = q(x_*) = y_0 + \frac{\frac{\partial g}{\partial x}(x_0, y_0)}{2\widehat{f}(x_0, y_0)}\delta^2 + \cdots .$$

Using $g(x_0, y_0)$ and the definition of $p_A$ in (33), we see that:

$$A_2(y_0) = -\frac{\frac{\partial g}{\partial x}(A - y_0, y_0)}{2p_A(y_0)}. \tag{47}$$

Similarly, by considering $\Psi_\delta(y)$, we find that:

$$B_2(y_0) = -\frac{\frac{\partial g}{\partial x}(A - y_0, y_0)}{2q_A(y_0)}. \tag{48}$$

Using (33) and (34), we have:

$$\begin{aligned} \frac{d}{dy}(A_1 + B_1)\bigg|_{y=y_0} &= \frac{d}{dy}\left(g(A - y, y)\left(\frac{1}{p_A(y)} + \frac{1}{q_A(y)}\right)\right)\bigg|_{y=y_0} \\ &= \frac{d}{dy}g(A - y, y)\bigg|_{y=y_0}\left(\frac{1}{p_A(y_0)} + \frac{1}{q_A(y_0)}\right), \end{aligned}$$

where we used $g(A - y_0, y_0) = 0$. Using the above together with (47), (48) and (45), we obtain (42). ☐

It is interesting that, in (42), the term linear in $\delta$ depends only on $g$ and its derivatives. In fact, using a geometric argument which we omit, it is possible to show that $y_\delta$ is approximated to order $\delta$ by the solution to the equation:

$$g(A - \widehat{y}_\delta - \delta/2, \widehat{y}_\delta) = 0.$$

It is easily checked that the expansion of $\widehat{y}_\delta$ with respect to $\delta$ agrees to first order in $\delta$ with that of $y_\delta$.

The following proposition states that, if there is an invariant region of (18), then there is a corresponding strict control region for $\mathcal{F}_\delta$ if $\delta$ is small enough.

**Proposition 2.5.** *Suppose $a \le y \le b, a < b$ is contained in the control region and*

$$g(A - a, a) = g(A - b, b) = 0,\ \frac{d}{dy}g(A - y, y)\bigg|_{y=a} \neq 0,\ \frac{d}{dy}g(A - y, y)\bigg|_{y=b} \neq 0.$$

*Then, there is a $\delta_* > 0$ such that for all $|\delta| < \delta_*$, there are values $a_\delta$ and $b_\delta$ satisfying $\mathcal{F}_\delta(a_\delta) = a_\delta$, $\mathcal{F}_\delta(b_\delta) = b_\delta$ with $a_{\delta=0} = a$ and $b_{\delta=0} = b$. Furthermore, $a_\delta < y < b_\delta$ is a strict control region for $\mathcal{F}_\delta$.*

*Proof.* That there is an $a_\delta$ satisfying $\mathcal{F}_\delta(a_\delta) = a_\delta,\ a_{\delta=0} = a_0$ is a direct consequence of item 1. of Proposition 2.4. The same holds for $b_\delta$. Take constants $c, d$ satisfying $c < a_\delta, b_\delta < d$ for all $|\delta| \le \delta_*$ such that $c \le y \le d$ resides within the control region. Note that:

$$\lim_{\delta \to 0} \frac{d\mathcal{F}_\delta}{dy} = 1 \text{ uniformly for } c \le y \le d.$$

Thus, by making $\delta_*$ smaller if necessary, we see that $\mathcal{F}_\delta$ is a strictly increasing function of $y$ for $a_\delta \leq y \leq b_\delta$ for $|\delta| \leq \delta_*$. Thus, $a_\delta < y < b_\delta$ implies:

$$a_\delta = \mathcal{F}_\delta(a_\delta) < \mathcal{F}_\delta(y) < \mathcal{F}(b_\delta) = b_\delta.$$

This shows that $a_\delta < y < b_\delta$ is a positively invariant region of $\mathcal{F}_\delta$. $\square$

The above results are of a somewhat general nature. Indeed, the proofs do not depend on the form of the function $g(x, y)$ in (5). In the corollary below, we apply the above results specifically to the function $g$ given in (7).

**Corollary 2.6.** *Suppose* $\alpha < 1, \beta > 1, A < 1, \beta A > 1$ *where* $\alpha, \beta$ *are as in* (6) *and* (7). *Suppose furthermore that:*

$$r_d > \widehat{R}_c(A - y_0^*, y_0^*), \; y_0^* = \frac{\beta A - 1}{\beta - 1}. \tag{49}$$

*Then, for* $\delta > 0$ *small enough, there is an unstable fixed point* $y_\delta^*$ *of* $\mathcal{F}_\delta$ *satisfying*

$$y_\delta^* = \frac{\beta A - 1}{\beta - 1} - \frac{\beta}{2(\beta - 1)}\delta + \mathcal{O}(\delta^2). \tag{50}$$

*Furthermore, suppose that:*

$$r_d > \widehat{R}_c(A - y, y) \text{ for } 0 \leq y \leq y_0^*. \tag{51}$$

*Then, for small enough* $\delta > 0$*, the open interval* $0 < y < y_\delta^*$ *is a strict control region. Within this strict control region,* $\widehat{R}_c$ *is in the anti-proliferative range:*

$$\widehat{R}_c(x, y) < r_x. \tag{52}$$

*Proof.* The above follows directly as an application of Theorem (2.4) and Proposition (2.5). The conditions $\beta > 1, A < 1, \beta A > 1$ ensures that the $y$ nullcline intersects with the line $x + y = A$ at the point $(A - y_0^*, y_0^*)$ and that $y_0^* > 0$ and $A - y_0^* > 0$. This implies in particular that $g(A - y_0^*, y_0^*) = 0$. Furthermore, we have:

$$\left.\frac{d}{dy} g(A - y, y)\right|_{y = y_0^*} = \beta A - 1 > 0.$$

Condition (49) ensures that the point $y_0^*$ is contained in the control region (see (36)). Thus, we may apply Theorem 2.4 to obtain the first half of the above statement. To prove the statement about the strict control region, we apply Proposition 2.5. Condition (51) ensures that $0 \leq y \leq y_0^*$ is in the control region. For any $\delta > 0$ and small enough, $y = 0$ is a fixed point of $\mathcal{F}_\delta$ as is easily seen by the fact that $g(x, y = 0) = 0$. The result follows. To see that (52) holds, note that $g(x, y) < 0$ in the strict control region. From this, we see that:

$$\widehat{R}_c(x, y) = \frac{f(x, y) + g(x, y)}{x} < \frac{f(x, y)}{x} = r_x(1 - x - \alpha y) \leq r_x.$$

$\square$

It can also be shown that, within the strict control region $0 < y < y_\delta^*$, $\mathcal{F}_\delta$ there are no fixed points, and that all points asymptotically approach $y = 0$. Thus, the resistant population goes extinct in this case. We omit this proof.

An interesting conclusion from the above is that, for $\delta > 0$, the strict control region for which the resistant population will go extinct is strictly smaller than the case of continuous adaptive therapy. This comes from the fact that $\frac{\partial g}{\partial x} < 0$, which is in turn a consequence of our assumption that the sensitive and resistant cell population are in competition.

#### 2.5 Intermittent Adaptive Therapy converges to Continuous Adaptive Therapy as $\delta \to 0$

We are now ready to address the comment at the end of Section 2.2. We will show that (18) provides a good approximation to the dynamics of intermittent adaptive therapy when $\delta$ is small enough. In other words, intermittent adaptive therapy converges to continuous adaptive therapy as $\delta \to 0$. An interesting technical point about the proof is that we work with a rescaled time variable $s$, rather than $t$. In this rescaled time $s$, the interval between rescaled switching times $s_{k+1} - s_k$ is approximately equal to $\delta$.

**Theorem 2.7.** *Consider the solution $z(t)$ of (18) with $z(0) = z_0$. Suppose the solution is defined for $0 \leq t \leq T$ and the set $\mathcal{S} = \{y = z(t) | 0 \leq t \leq T\}$ is in the control region. Consider the differential equations of intermittent adaptive therapy with $A$ and $A - \delta$ be the upper and lower thresholds of tumor size, and let $x = x_\delta(t), y = y_\delta(t)$ be the solution to this system with initial data $x_\delta(0) = A - z_0$ and $y_\delta(0) = z_0$. Then, there is a $\delta_* > 0$ such that, for all $|\delta| < \delta_*$ we have:*

$$A - \delta \leq x_\delta(t) + y_\delta(t) \leq A \text{ for all } 0 \leq t \leq T \tag{53}$$

*and*

$$\sup_{0 \leq t \leq T} |y_\delta(t) - z(t)| \leq C\delta \tag{54}$$

*where the constant $C$ only depends on $T$.*

*Proof.* Consider the following system of ordinary differential equations:

$$\frac{dw}{ds} = P(w),\ P(w) = g(A - w, w) \left( \frac{1}{p_A(w)} + \frac{1}{q_A(w)} \right), \tag{55}$$

$$\frac{d\tau}{ds} = Q(w),\ Q(w) = \frac{1}{p_A(w)} + \frac{1}{q_A(w)},\ w(0) = z_0,\ \tau(0) = 0. \tag{56}$$

First, note that the above differential equation and (18) are equivalent by the assumption that $\mathcal{S}$ is in the control region. Indeed for any $y \in \mathcal{S}$, we know that $Q(y)$ is positive and bounded, and thus, we may use $s$ above as our new "time" variable. That is to say, $w = z(\tau(s))$ for $0 \leq s \leq S$ where $\tau(S) = T$.

We may extend this solution further still to $S_* > S$. We choose $S_*$ so that $\mathcal{S}_* = \{y = z(t) | 0 \leq t \leq T_* = \tau(S_*)\}$ is still contained in the control region. Note that, since $Q(y) > 0$ for $y \in \mathcal{S}_*$, $\tau$ is monotone increasing and $T_* = \tau(S_*) > \tau(S) = T$. Let $\mathcal{S}_* = \{a \leq y \leq b\}$. That is to say, either $a = z(0)$ or $a = z(T_*)$ and similarly for $b$. We may pick a $c > 0$ so that the slightly expanded interval $\mathcal{S}_{**} = \{a - c \leq y \leq b + c\}$ is also still in the control region.

For $\delta > 0$, pick the largest integer $N_\delta$ that satisfies $N_\delta \delta \leq S_*$. Let $s_k = k\delta$ and $w_k = w(s_k), \tau_k = \tau(s_k)$. Note that $w_k \in \mathcal{S}_*$. We have

$$w_{k+1} = w_k + P(w_k)\delta + R_w(w_k, \delta), \tag{57}$$

$$\tau_{k+1} = \tau_k + Q(w_k)\delta + R_\tau(w_k, \delta). \tag{58}$$

The remainder terms $R_w$ and $R_\tau$ satisfy the estimates

$$|R_w(w, \delta)| \leq K_1 \delta^2,\ |R_\tau(w, \delta)| \leq K_1 \delta^2$$

where $K_1$ does not depend on $|\delta| < \delta_*$ for some $\delta_* > 0$ and $w \in \mathcal{S}_*$.

Let $v_0 = z_0, \sigma_0 = 0$ and define:

$$y_{k+1} = \mathcal{F}_\delta(y_k),\ t_{k+1} = t_k + \mathcal{T}_\delta(t_k). \tag{59}$$

Note that $\mathcal{F}_\delta, \mathcal{T}_\delta$ are defined so long as $y_k \in \mathcal{S}_{**}$ and $|\delta| < \delta_*$, by making $\delta_*$ smaller (but positive) if necessary. The values $t_k$ are precisely the switching times for intermittent adaptive therapy (see (16)), and $y_k = y_\delta(t_k)$. Using Lemma 2.3, we see that:

$$y_{k+1} = y_k + P(y_k)\delta + R_y(y_k, \delta), \tag{60}$$

$$t_{k+1} = t_k + Q(y_k)\delta + R_t(y_k, \delta). \tag{61}$$

The remainder terms $R_y$ and $R_t$ satisfy the estimates

$$|R_y(y, \delta)| \leq K_2\delta^2,\ |R_t(y, \delta)| \leq K_2\delta^2$$

where $K_2$ does not depend on $|\delta| < \delta_*$ (by making $\delta_*$ smaller if necessary, but positive) and $y \in \mathcal{S}_{**}$. Let $w_k - y_k = e_k$, so long as $w_k$ and $y_k$ are defined. Suppose $w_k \in \mathcal{S}_*$ and $y_k \in \mathcal{S}_{**}$ and $|\delta| < \delta_*$. Using (57) and (60), we have:

$$|e_{k+1}| \leq (1 + \delta L)\,|e_k| + K_3\delta^2,\ e_0 = 0,$$

where $L$ is the Lipschitz constant of $P$ in $\mathcal{S}_{**}$ and $K_3 = K_1 + K_2$. We may conclude from the above inequality that, so long as $w_\ell \in \mathcal{S}_*$ and $y_\ell \in \mathcal{S}_{**}$ for all $0 \leq \ell \leq k-1$, we have:

$$|e_k| \leq \frac{K_3\delta}{L}\left((1+\delta L)^k - 1\right).$$

Now, note that, if $k \leq N_\delta$,

$$\frac{K_3\delta}{L}\left((1+\delta L)^k - 1\right) \leq \frac{K_3\delta}{L}\left(\exp(LS_*) - 1\right).$$

By taking $\delta_*$ smaller if necessary so that:

$$\frac{K_3\delta_*}{L}\left(\exp(LS_*) - 1\right) < c$$

we see that $|e_{k+1}| < c$ so long as $w_\ell \in \mathcal{S}_*$ and $y_\ell \in \mathcal{S}_{**}$ for all $0 \leq \ell \leq k-1$. Note that, $0 \leq \ell \leq N_\delta$, $w_\ell \in \mathcal{S}_*$. By the definition of $\mathcal{S}_{**}$, we recursively conclude that $y_\ell \in \mathcal{S}_{**}$ for $0 \leq \ell \leq N_\delta$. Thus, for $0 \leq k \leq N_\delta$, we have:

$$|w_k - y_k| \leq \frac{K_3\delta}{L}\left(\exp(LS_*) - 1\right) \text{ for } |\delta| < \delta_*.$$

Using (58) and (61), the Lipschitz continuity of $Q$ and the above estimate, we find:

$$|\tau_k - t_k| \leq M\delta \text{ for } |\delta| < \delta_*,$$

where $M$ depends only on $S_*$. This shows in particular that:

$$t_{N_\delta} \geq \tau_{N_\delta} - M\delta \geq \tau(S_* - \delta) - M\delta \text{ for } |\delta| < \delta_*.$$

Since $\tau(S_* - \delta) \to \tau(S_*) = T_*$ as $\delta \to 0$, we can make $\delta_*$ smaller if necessary so that $\tau(S_* - \delta) - M\delta > T$. Thus, the sequence of switching times $0 = t_0 < t_1 < \cdots < t_{N_\delta}$ extends beyond $T$ for $|\delta| < \delta_*$. By the definition of switching times, we see that (53) holds. The bound (54) is now an immediate consequence of (53) and Proposition 2.1. □

#### 2.6 Time is Optimized with Continuous Adaptive Therapy

In this section, the time to move the population along the control line to either a higher or lower value of resistant species, $y$, is examined. We show that time to resistant species reduction is minimized under continuous control (or equivalently, as $\delta \to 0$) and time to resistant species outgrowth is maximized under continuous control.

**Proposition 2.8.** *Let $z(t), x_\delta(t), y_\delta(t)$, $0 \le t \le T < \infty$ satisfy the same assumptions of Theorem 2.7. Assume that $z'(0) > 0$. This implies that $z'(t) > 0$ for $0 \le t \le T < \infty$. Suppose that:*

$$\frac{\partial g}{\partial x}(A - z, z) < 0 \text{ for } z(0) = z_0 \le z \le z_1 = z(T). \tag{62}$$

*Then, there is a $\delta_* > 0$ such that for all $0 < \delta \le \delta_*$, there is a $T_\delta$ such that $y_\delta(T_\delta) = z_1$ and $y'_\delta(t) > 0$ for $0 \le t \le T_\delta$. Furthermore, $T_\delta < T$.*

The time $T$ is the time to resistant species outgrowth for continuous adaptive therapy whereas $T_\delta$ is the corresponding time for intermittent adaptive therapy. Note that this result depends crucially on (64), which says that there is a competition between sensitive and resistant species. Indeed,

$$\frac{\partial g}{\partial x} = -r_y \beta y$$

and thus, as long as $y$ is positive, this is always negative. The proof below depends on $\delta$ being small, but the result that $T_\delta < T$ is likely to hold for a much wider range of conditions.

*Proof of Proposition 2.8.* That $z'(0)$ implies $z'(t) > 0$ for $0 \le t \le T$ follows from the fact that (18) is a one-dimensional differential equation in which $g(A - z, z)$ is a smooth function. The assertion that, $y_\delta(t)$ exists up until $y_\delta$ reaches $z_1$ follows from Theorem 2.7. Furthermore, we can prove that $x_\delta, y_\delta$ is in the weak control region and consequently

$$A - \delta \le x_\delta(t) + y_\delta(t) \le A \text{ for } 0 \le t \le T_\delta, \quad |T_\delta - T| \le C\delta \tag{63}$$

where $C$ does not depend on $\delta$. We omit the details of this proof. Note that:

$$g(A - y, y) > 0, \text{ for } z_0 \le y \le z_1.$$

Since $A - \delta - y_\delta \le x_\delta \le A - y_\delta$, we see that, for $\delta$ small enough,

$$g(x_\delta(t), y_\delta(t)) > 0, \text{ for } 0 \le t \le T_\delta.$$

This implies that $y_\delta(t)$ is monotone increasing. This then implies that we can write $x_\delta(t)$ as a function of $y_\delta(t)$:

$$x_\delta = X_\delta(y_\delta(t)).$$

From this, we see that:

$$T_\delta = \int_{z_0}^{z_1} \frac{1}{g(X_\delta(z), z)} dz.$$

On the other hand,

$$T = \int_{z_0}^{z_1} \frac{1}{g(A - z, z)} dz.$$

Note that $X_\delta(z) \le A - z$. Given condition (64), $g(X_\delta(z), z) \ge g(A - z, z)$ for $\delta$ small enough. Thus, $T_\delta \le T$. In fact, $T_\delta < T$ since $X_\delta(z)$ is not identically equal to $A - z$. □

We have the following result for resistance species extinction.

**Proposition 2.9.** *Let* $z(t), x_\delta(t), y_\delta(t)),\ 0 \le t \le T < \infty$ *satisfy the same assumptions of Theorem 2.7. Assume that* $z'(0) < 0$*. This implies that* $z'(t) < 0$ *for* $0 \le t \le T < \infty$*. Suppose that:*

$$\frac{\partial g}{\partial x}(A - z, z) < 0 \text{ for } z(T) = z_1 \le z \le z_0 = z(0). \tag{64}$$

*Then, there is a* $\delta_* > 0$ *such that for all* $0 < \delta \le \delta_*$*, there is a* $T_\delta$ *such that* $y_\delta(T_\delta) = z_1$ *and* $y'_\delta(t) > 0$ *for* $0 \le t \le T_\delta$*. Furthermore,* $T_\delta > T$*.*

The proof of this result is almost exactly the same as Proposition 2.8.

A simulation of this result is shown in Figure 14 and demonstrates that as $\delta$ decreases, the time to disease control also decreases.

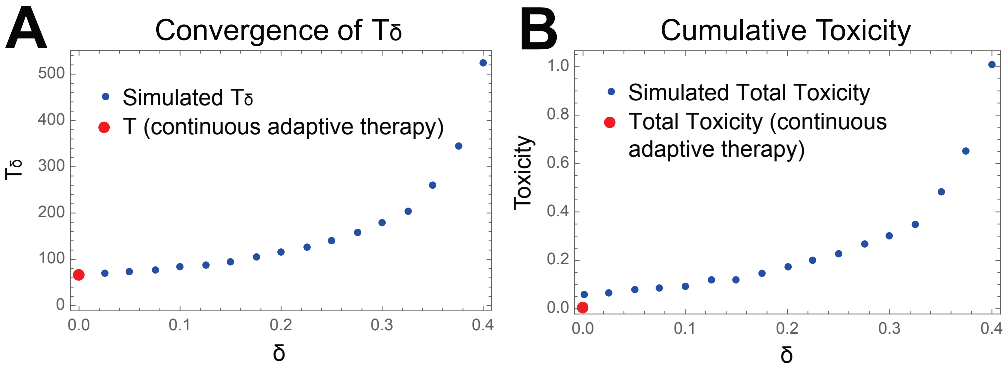

Figure 14: **Time to disease control and cumulative toxicity**. **A** Numerical simulation of the time $T_\delta$ to reach $y = \epsilon$. It can be seen that $T_\delta$ converges to $T$(see Proposition 2.8) as $\delta \to 0$. **B** Numerical simulation of the cumulative toxicity for the simulation shown in A. Toxicity is modeled as $r_d^2 t$. Total toxicity for continuous adaptive therapy is shown in red and is strictly less than drug toxicity in intermittent adaptive therapy (see Proposition 2.11.) *Simulation Parameters:* $A = 0.8$, $a = 0.5$, $b = 2.1$, $x_0 = 0.6$, $y_0 = 0.2$, $r_d = 0.1$, $r_x = 0.04$, $r_y = 0.02$, *and Cumulative Toxicity* $= r_d^2 t$.

#### 2.7 Continuous Adaptive Therapy and Drug Toxicity

Consider intermittent adaptive therapy with drug level $r_d$. Given drug level $r_d > 0$, suppose $D(r_d)$ is the level of toxicity. Consider the cumulative toxicity up to time $T$:

$$\mathcal{D}_\delta(T) = D(r_d)\Theta_\delta(T, r_d),\ \Theta_\delta(T, r_d) = \int_0^T K_A(t)dt.$$

This is just the product of $D(r_d)$ with $\Theta_\delta(T, r_d)$, the length of time over which the drug was turned on. Note that $K_A(t)$ depends on $r_d$ and $\delta$. Let us consider cumulative drug toxicity, where at $t = 0$, $x_\delta(0) + y_\delta(0) = A$.

**Proposition 2.10.** *Under the same assumption as in Theorem 2.7, we have:*

$$\lim_{\delta\to 0}\mathcal{D}_\delta(T) = D(r_d)\int_0^T \frac{q_A(z(t))}{r_d(A-z(t))}dt.$$

*Proof.* We shall use the same notation as the proof of Proposition 2.7. Let $v_k = \Theta_\delta(t_k, r_d)$ be the cumulated time over which drug was turned on up till $t = t_k$. Then,

$$v_{k+1} = v_k + \mathcal{T}_\delta^\Phi(y_k).$$

From the proof of Lemma 2.3, we have:

$$v_{k+1} = v_k + \frac{1}{p_A(y_k)}\delta + R_v(y_k,\delta),\ |R_v(y,\delta)| \le K\delta^2,$$

where $K$ independent of $|\delta| < \delta_*$ and $y \in \mathcal{S}_*$.

Augment the differential equations (55) and (56) with the differential equation:

$$\frac{du}{ds} = \frac{1}{p_A(w)},\ u(0) = 0.$$

Using the same argument as in the proof of Proposition 2.7, we see that:

$$|v_k - u(s_k)| \le M_1\delta \text{ for } |\delta| < \delta_*,\ 0 \le k \le N_\delta$$

where $M_1$ is a constant that is independent of $\delta$. For each $\delta$, let $k_\delta$ be the largest integer such that $s_{k_\delta} = k_\delta\delta \le S$, so that $|S - s_{k_\delta}| < \delta$. Then, we have:

$$|v_{k_\delta} - u(s_{k_\delta})| \le M_1\delta.$$

Note that:

$$\begin{aligned}|v_{k_\delta} - \Theta_\delta(T,r_d)| &= |\Theta_\delta(t_{k_\delta}, r_d) - \Theta_\delta(T, r_d)| \le |T - \tau(s_{k_\delta})|\\ |\Theta_\delta(T,r_d) - u(S)| &\le |\Theta_\delta(T,r_d) - v_{k_\delta}| + |v_{k_\delta} - u(s_{k_\delta})| + |u(s_{k_\delta}) - u(S)|\\ &\le |T - \tau(s_{k_\delta})| + M_1\delta + \int_{s_{k_\delta}}^S \frac{1}{p(w(s))}ds.\end{aligned}$$

Letting $\delta \to 0$, we see that:

$$\lim_{\delta\to 0}\Theta_\delta(T, r_d) = u(S) = \int_0^S \frac{1}{p_A(w(s))}ds.$$

Using the relation $\tau(0) = 0, T = \tau(S), z(\tau(s)) = w(s)$ and the fact that $\tau$ is monotone increasing, we may change the variable of integration to obtain:

$$\int_0^S \frac{1}{p_A(w(s))}ds = \int_0^T \frac{q_A(z(t))}{p_A(z(t)) + q_A(z(t))}dt.$$

We obtain the desired result by noting that $p_A(z) + q_A(z) = -h(A-y, r_d) = r_d(A-y)$ (see (33) and (34)). □

Let us use the above result to compare the cumulative drug toxicity for intermittent adaptive therapy with that of continuous adaptive therapy. For continuous adaptive therapy, drug toxicity is given by:

$$\mathcal{D}_{\text{cont}} = \int_0^T D(R_c(A - z(t), z(t)))dt = \int_0^T D\left(\frac{q_A(z(t))}{A - z(t)}\right) dt. \tag{65}$$

where we used (30), (34) as well as the fact that $x = A - z(t), y = z(t)$ for continuous adaptive therapy.

Now, let us make some assumptions on the function $D$ relating drug level $r_d$ to toxicity. We assume the following for $D$.

1. $D(w)$ is continuous function defined for $w \geq 0$ and satisfies $D \geq 0$.

2. For any $0 < \alpha < 1$ and $w \geq 0$ we have $D(\alpha w) < \alpha D(w)$.

One class of functions $D$ that satisfy the above conditions are those that satisfy $D(0) = 0, D' > 0, D'' > 0$. We have the following result.

**Proposition 2.11.** *Suppose the drug toxicity function $D$ satisfy the properties above. Under the same assumptions as in Theorem 2.7,*

$$\lim_{\delta \to 0} \mathcal{D}_\delta(T) > \mathcal{D}_{\text{cont}}(T).$$

The point here is that the limit of cumulative toxicity as $\delta \to 0$ is strictly larger than that of continuous adaptive therapy. This result is true for a wider class of conditions for $D$; so long as $D(\alpha w) \leq \alpha D(w)$ is satisfied for $0 < \alpha < 1$ and $D(w)$ is not linear in $w$, then, this result is true so long as $r_d$ is large enough. From a mathematical point of view, this strict inequality comes from the fact that $r_d K_A(t)$ is converging to $R_c(A - z(t), z(t))$ only in a weak sense.

*Proof.* Since the trajectory resides in $\mathcal{C}$, the area of controllability (see (35)), we have from (36):

$$0 < \frac{R_c(A - z(t), z(t))}{r_d} = \frac{q_A(z(t))}{r_d(A - z(t))} < 1.$$

Thus,

$$D(r_d)\frac{q_A(z(t))}{r_d(A - z(t))} > D\left(\frac{q_A(z(t))}{A - z(t)}\right).$$

The result follows from Proposition 2.10 and (65). □

A simulation of the above result is shown in Figure 14B.

### 3 Population Dynamics under Continuous Fixed Dose Treatment

In this section, the dose effect relationship of continuous administration of chemotherapy in the modified Lotka-Volterra competition model is examined in order to determine optimal doses for continuous fixed-dose therapy. It is demonstrated that this optimal dosage is in the antiproliferative range of drug effect and that there are certain concentrations of continuous fixed dose antiproliferative drug that result in resistant species extinction when $\alpha < 1$ and $\beta > 1$.

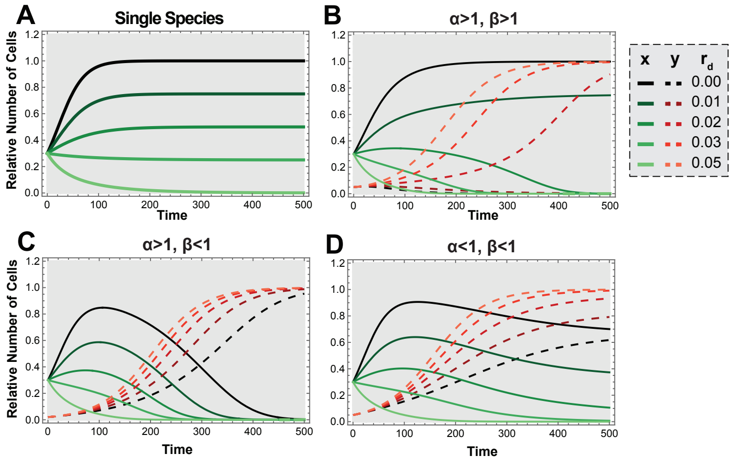

Figure 15: **Dose effect of drug treatment under varying competition parameters. (A)** Single species treatment for given drug doses ($r_d$) with $x_0 = 0.3$. **(B)** $\alpha = 2$, $\beta = 2$, and $(x_0, y_0) = (0.3, 0.05)$. **(C)** $\alpha = 0.5$, $\beta = 0.5$, and $(x_0, y_0) = (0.3, 0.05)$. **(D)** $\alpha = 2$, $\beta = 0.5$, $(x_0, y_0) = (0.3, 0.02)$. *** Other simulation parameters:* $r_x = 0.04$ *and* $r_y = 0.02$

#### 3.1 Dose-effect Relationship in Competing Cell Populations

In the main text, single species behavior is simulated over time in Figure 5A for a range of drug concentrations spanning cytotoxic ($r_d > r_x$) to antiproliferative ($r_d < r_x$). It is readily observed that the carrying capacity changes depending on the dose of drug. Given a desired equilibrium, $x_{eq}$ for $x$, Equation 12 can be rearranged to yield:

$$r_d = r_x(1 - x_{eq}) \tag{66}$$

in order to set a new carrying capacity at any given value.

When a second species, $y$, is added, the effectiveness of the drug in reducing the population of $x$ is amplified further. Figure 5B shows the case where the sensitive cells outcompete the resistant cells in the absence of drug. For drug concentrations that were previously antiproliferative rather than cytotoxic, the decline in the sensitive cell population allows the resistant population to compete and effectively drive the sensitive cell population to extinction.

In the case where the resistant cell population is a stronger competitor than the sensitive cell population, no continuous treatment strategy can result in resistant cell population eradication, however, the time to progression shortens as the drug dose increases (Supplemental Figure 15C). Conceptually, this occurs because the higher the tumor volume (which is predominantly sensitive cells at first), the lower the growth rate of the resistant cell population. The implication of this observation is that in systems where there is no strategy that results in resistant cell population control, lower doses of continuous drug result in a longer duration of time before the resistant population overtakes the sensitive population. Additional simulations are shown in 15B-C).

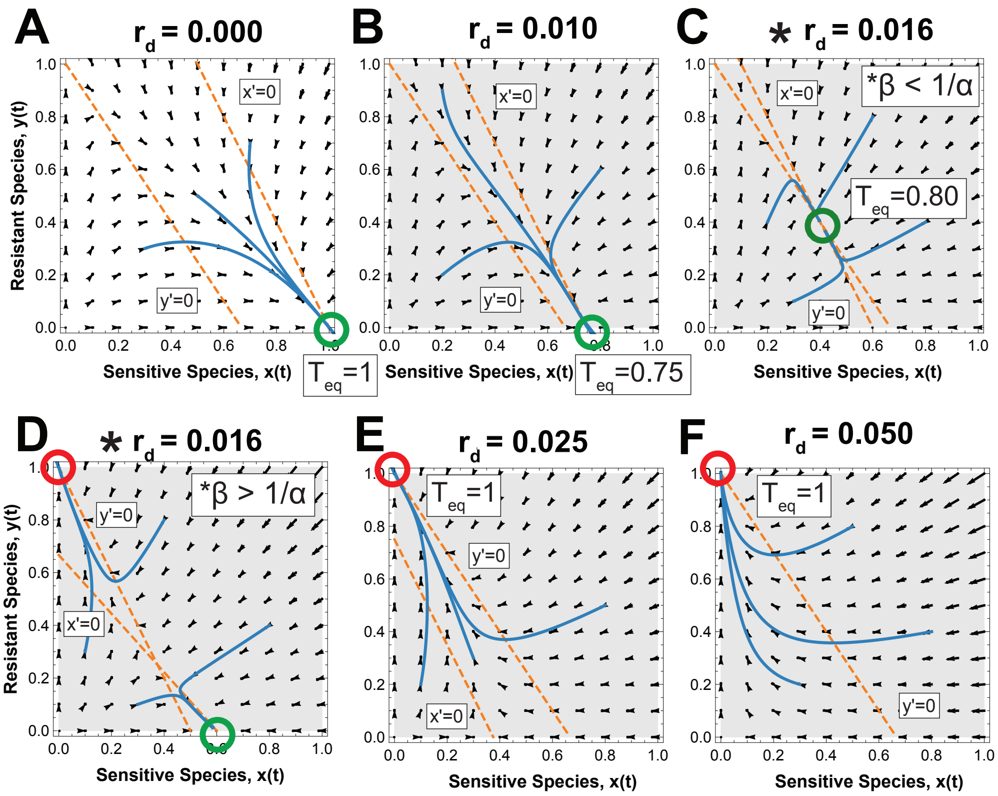

Figure 16: **Phase planes for varying continuous fixed-dose dose therapy when $\alpha < 1$ and $\beta > 1$.** As $r_d$ varies, the relationship of the $x$ and $y$ nullclines changes the observed equilibrium points. Blue lines indicate trajectories in phase space for to different initial conditions. (**C**) The intersection of the $x$ and $y$ nullclines when $\beta < \frac{1}{\alpha}$ is a stable equilibrium representing a mixed tumor population. (**D**) The intersection of the $x$ and $y$ nullclines when $\beta > \frac{1}{\alpha}$ creates two stable equilibria yielding either completely resistant or completely sensitive cell populations. *Simulation Parameters:* $r_x = 0.04$, $r_y = 0.03$. $\alpha = 0.5$ $\beta = 1.5$ for A, B, C, E, and F. $\alpha = 0.9$ $\beta = 2$ for D.

#### 3.2 Optimal Continuous Fixed Dose Therapy when $\alpha < 1$ and $\beta > 1$

When the sensitive population can outcompete the resistant population, optimal dosing strategies to achieve control for a defined maximal tumor volume, $A$, can be directly explored.

The nullclines under continuous dosing ($K_A = 1$) for this system are given below

$$y = \frac{1 - x - \frac{r_d}{r_x}}{\alpha} \qquad x\text{-nullcline} \tag{67}$$

$$y = 1 - \beta x \qquad y\text{-nullcline} \tag{68}$$

and are plotted in Figure 5 with the corresponding phase planes in Supplementary Figure 16 for

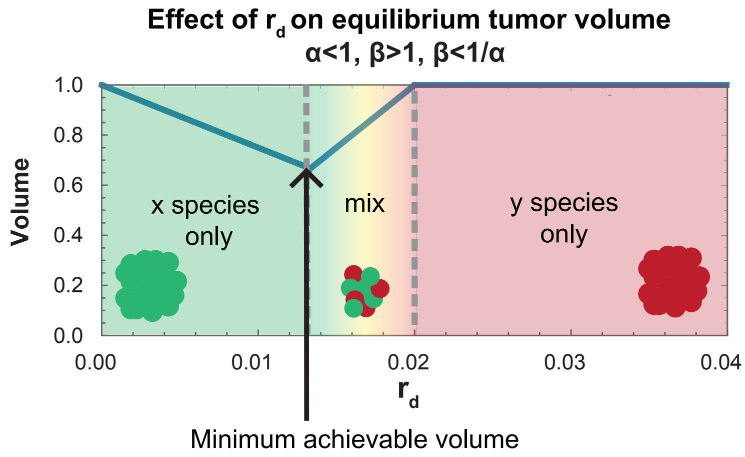

Figure 17: **Cartoon depiction of the minimum tumor volume.**

varying values of $r_d$. Note that these values of $r_d$ and parameters correspond to those shown for the time course in Figure 5A and B.

The effect of drug in this system is to push the $x$-nullcline toward 0, creating a series of values of $r_d$ for which the $x$ and $y$ nullclines intersect. When $\beta < \frac{1}{\alpha}$ this intersection forms a stable equilibrium where the sensitive and resistant species coexist Figure 16C. When $\beta > \frac{1}{\alpha}$, this equilibrium is unstable and the system tends to completely resistant or completely sensitive (Supplementary Figure 16D).

We can now solve for the minimum achievable stable tumor size using continuous dosing in this system using phase planes.

**Proposition 3.1.** *For $\alpha < 1$, $\beta > 1$, and $\beta < \frac{1}{\alpha}$, the minimum achievable tumor size with continuous treatment is $\frac{1}{\beta}$ for any initial conditions.*

*Proof.* By phase plane argument, the the minimum equilibrium tumor volume achievable by varying $r_d$ occurs when the $x$-nullcline intersects the $y$-nullcline on the $x$ axis. This intersection occurs at $x = \frac{1}{\beta}$ and $(\frac{1}{\beta}, 0)$ becomes the only stable equilibrium. □

**Corollary 3.2.** *The dose, $r_d$, needed to achieve this minimum tumor volume given in Proposition 3.1 is now explicitly solvable and yields:*

$$r_d = \frac{r_x(\beta - 1)}{\beta} \tag{69}$$

A cartoon representation of the minimum tumor volume is shown in Figure 17. In biological terms, if the maximum tolerated tumor volume, $A$, is greater than $\frac{1}{\beta}$, then there is a dose $r_d$ that allows for extinction of the resistant population with the equilibrium tumor volume less than $A$ for any initial conditions.

In the case where $\beta > \frac{1}{\alpha}$, the minimum achievable tumor volume is dependent on initial conditions. When the initial conditions allow the trajectory to reach the $x$ only equilibrium, then the minimum tumor volume becomes the intersection of the $x$-nullcline with the $x$-axis.

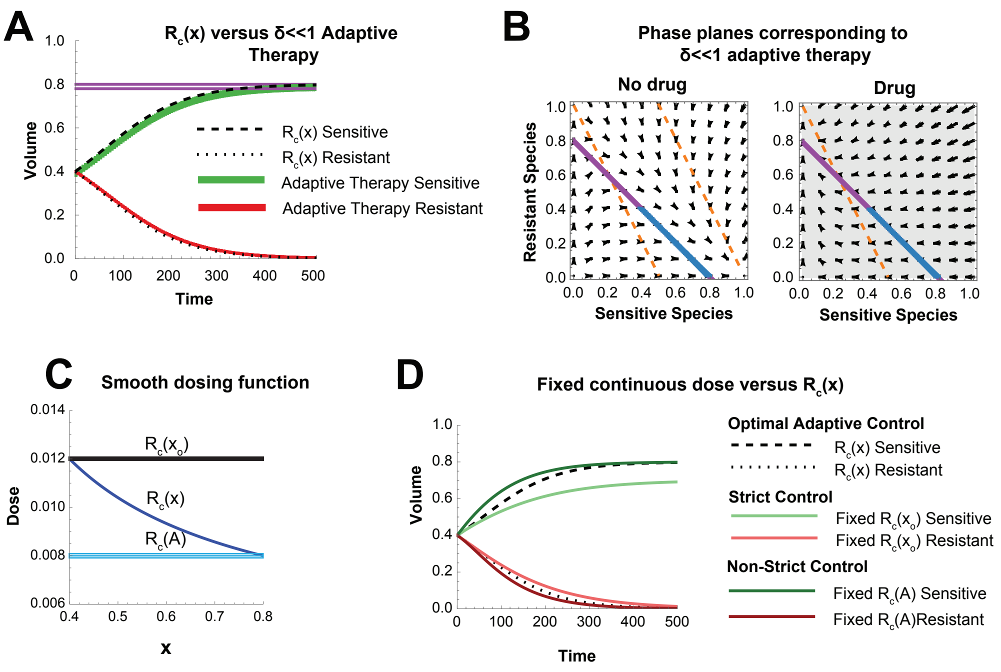

Figure 18: **Direct comparison of optimal ($\delta \ll 1$) adaptive therapy versus optimal fixed dose therapy.** **(A)** Comparison of $\delta \ll 1$ adaptive therapy (green and red trajectories with control lines shown in purple) versus the two species model with the continuous dosing function $R_c(x)$ representing $\delta = 0$. **(B)** Corresponding phase plane for $\delta \ll 1$ shown in Figure (A). **(C)** Smooth dosing function $R_c(x)$ with upper and lower bounds $R_c(x_0)$ and $R_c(A)$ plotted. **(D)** System behavior with $R_c(x)$ (dotted lines) versus continuous fixed-dose therapy with dose fixed at $R_c(x_0)$ or $R_c(A)$.

#### 3.3 Resistant Population Extinction under Fixed-Dose Continuous Therapy versus Adaptive Therapy for $\alpha < 1$ and $\beta > 1$

Results of a simulation of the two species model with continuous versus intermittent adaptive therapy with $\delta \ll 1$ are shown in Figure 18A and demonstrate excellent concordance. The phase planes corresponding to $\delta \ll 1$ are shown in Figure 18B with the unstable limit cycle, strict control region, and control region as described previously.

In order to directly compare continuous adaptive therapy with continuous fixed dose therapy, function $R_c(x) = \widehat{R}_c(x, A - x)$ (see Eq. (30)) is plotted in a segment of the strict control region (Figure 18C). Tumor control can be achieved in this case at the lowest dose ($R_c(x) = R_c(A)$) and the highest dose ($R_c(x_0)$) given continuously at a fixed concentration. The resulting simulation is shown in Figure (Figure 18D).

For these parameters, it is noted that restriction of the final tumor volume to A with some growth allowance above $A$ for a finite period of time can be achieved with a low fixed anti-proliferative dose of $R_c(x)$. The time to resistant extinction, in this case, is shorter than optimal adaptive therapy. Under strict tumor size control, the higher continuous fixed dose treatment results in tumor volume at or below $A$ for the duration of therapy. In this case, the time to resistant extinction is larger relative to optimal adaptive therapy, however, the steady state tumor volume is smaller relative to optimal adaptive therapy

###### 3.3.1 $\beta < \frac{1}{\alpha}$

We are now able to directly compare optimal adaptive therapy to optimal fixed dose therapy in terms of the relative sensitivity to initial conditions. When $\beta < \frac{1}{\alpha}$ and $A > \frac{1}{\beta}$, there exists an $r_d$ such that for any initial conditions, the resistant population can be driven to extinction (Proposition 3.1). In adaptive therapy, on the other hand, the separatrix shown in Figure 5 demonstrates that whether or not disease control is achievable under adaptive therapy is dependent on the initial conditions.

###### 3.3.2 $\beta > \frac{1}{\alpha}$

When $\beta > \frac{1}{\alpha}$, the relative positions of the nullclines (Figure 5D determine which therapy modality is less sensitive to initial conditions. Recall that, unlike $\beta < \frac{1}{\alpha}$, the success of continuous fixed dose therapy is dependent on initial conditions.

Assuming a the most important constraint is strict tumor size control below $A$, $r_d$ can be chosen such that the maximum tumor size is always less than $A$ by setting the intersection of the $x$ and $y$ nullclines on the $x + y = A$ line. Here, the fixed continuous antiproliferative drug dose would be applied when the tumor reaches the volume $A$ and held until resistant population extinction. The phase space comparison with adaptive therapy is shown in Figure 5. The initial conditions allowing for tumor control are equal between adaptive and continuous therapy, however, continuous antiproliferative therapy allows the maintenance of a lower tumor volume overall as the resistant population is being driven extinct.

If the tumor size constraint is non-strict, then the region of phase space for which continuous antiproliferative therapy allows for resistant population extinction is larger than that of adaptive therapy (Figure 5).

Taken together, fixed antiproliferative dosing allows for extinction of the resistant population for the same or larger set of initial conditions as optimal adaptive therapy depending on the strictness of the tumor volume constraint.

##### 3.4 Comparison with Continuous Fixed Dosing when $\alpha > 1$ and $\beta < 1$

When the resistant population extinction cannot be achieved, direct comparison of the two modalities is less informative. A simulation is shown in Figure 19. This simulation demonstrates behavior in the phase space shown in Figure 13 where the $x + y = A$ line is in the strict control region for part of the trajectory and returns to the region of controllability for the second part of the trajectory. Again, excellent concordance between the continuous $R_c$ dosing function and $\delta \to 0$ adaptive therapy is shown (Figure 19A,B).

When plotting $R_c(x)$, we note that when the system is out of the region of controllability and below the $x + y = A$ line, $R_c(x)$ takes values below 0 (Figure 19C) and thus continuous adaptive therapy is not defined there. Applying a similar upper and lower boundary of $R_c$ approach as in Figure 18 allows for comparison of no treatment (non-strict tumor size control) with continuous fixed dose antiproliferative treatment ($r_d = R_c(x_0)$) (Figure 19D).

In this case, optimal adaptive therapy is superior to the higher fixed-dose antiproliferative therapy in terms of the time to resistant population progression. It is important to note, however, that the difference is small and that wider control is likely to negate this effect.

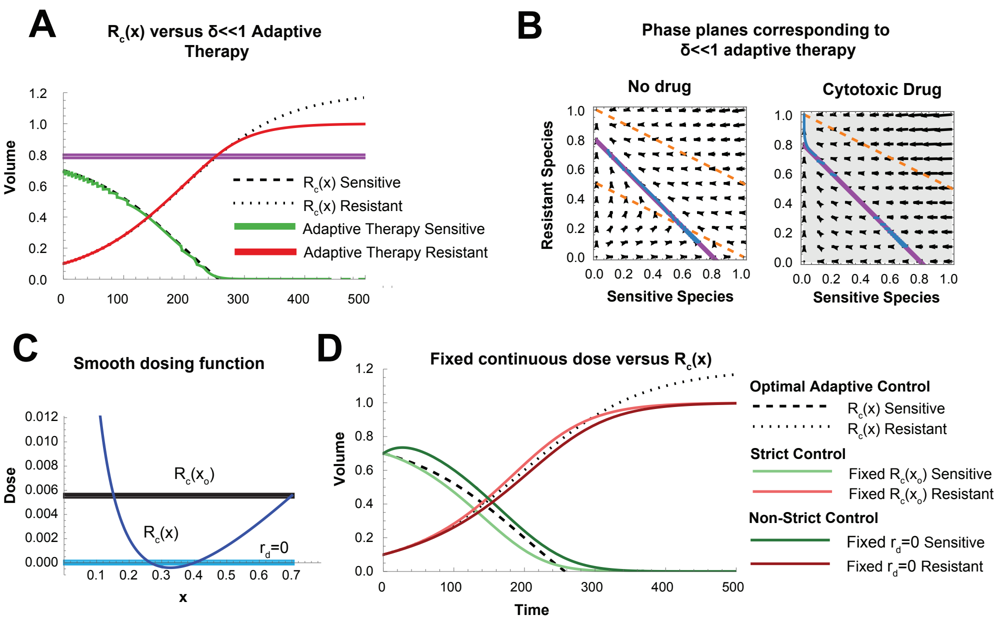

Figure 19: **Direct Comparison of $\delta \ll 1$ adaptive therapy versus fixed dose therapy (A)** Comparison of $\delta \ll 1$ adaptive therapy (green and red trajectories with control lines shown in purple) versus the two species model with the continuous dosing function $R_c(x)$ representing $\delta = 0$. **(B)** Corresponding phase plane for $\delta \ll 1$ shown in Figure (A). Note that the upper and lower control boundaries are not contained within the area of controllability for part of the trajectory. **(C)** Smooth dosing function $R_c(x)$ with fixed continuous dose comparison at $R_c(x_o)$ and $r_d = 0$ plotted. **(D)** System behavior with $R_c(x)$ (dotted lines) versus continuous fixed-dose therapy with dose fixed at $R_c(x_o)$ and $r_d = 0$.

### References

[1] Robert A Gatenby et al. "Adaptive therapy". In: *Cancer research* 69.11 (2009), pp. 4894–4903.

[2] Jingsong Zhang et al. "Integrating evolutionary dynamics into treatment of metastatic castrate-resistant prostate cancer". In: *Nature communications* 8.1 (2017), pp. 1–9.

[3] Nadia Howlader et al. "The effect of advances in lung-cancer treatment on population mortality". In: *New England Journal of Medicine* 383.7 (2020), pp. 640–649.

[4] Angela B Mariotto et al. "Estimation of the number of women living with metastatic breast cancer in the United States". In: *Cancer Epidemiology, Biomarkers & Prevention* 26.6 (2017), pp. 809–815.

[5] Ian Smith. "Goals of treatment for patients with metastatic breast cancer". In: *Seminars in oncology*. Vol. 33. Elsevier. 2006, pp. 2–5.

[6] SKIPPER HE. "Implications of biochemical, cytokinetics, pharmacologic, and toxicologic relationships in the design of optimal therapeutic schedules". In: *Cancer Chemother Rep* 54 (1970), pp. 431–450.

[7] Douglas Hanahan, Gabriele Bergers, Emily Bergsland, et al. "Less is more, regularly: metronomic dosing of cytotoxic drugs can target tumor angiogenesis in mice". In: *The Journal of clinical investigation* 105.8 (2000), pp. 1045–1047.

[8] Irina Kareva, David J Waxman, and Giannoula Lakka Klement. "Metronomic chemotherapy: an attractive alternative to maximum tolerated dose therapy that can activate anti-tumor immunity and minimize therapeutic resistance". In: *Cancer letters* 358.2 (2015), pp. 100–106.

[9] Nicolas André, Manon Carré, and Eddy Pasquier. "Metronomics: towards personalized chemotherapy?" In: *Nature reviews Clinical oncology* 11.7 (2014), pp. 413–431.

[10] Luc G T Morris et al. "Pan-cancer analysis of intratumor heterogeneity as a prognostic determinant of survival." eng. In: *Oncotarget* 7.9 (2016), pp. 10051–10063. ISSN: 1949-2553 (Electronic); 1949-2553 (Linking). DOI: 10.18632/oncotarget.7067.

[11] Jingsong Zhang et al. "Evolution-based mathematical models significantly prolong response to abiraterone in metastatic castrate-resistant prostate cancer and identify strategies to further improve outcomes". In: *eLife* 11 (2022). Ed. by George H Perry, e76284. ISSN: 2050-084X. DOI: 10.7554/eLife.76284. URL: https://doi.org/10.7554/eLife.76284.

[12] Pedro M Enriquez-Navas et al. "Exploiting evolutionary principles to prolong tumor control in preclinical models of breast cancer". In: *Science translational medicine* 8.327 (2016), 327ra24–327ra24.

[13] Inna Smalley et al. "Leveraging transcriptional dynamics to improve BRAF inhibitor responses in melanoma". In: *EBioMedicine* 48 (2019), pp. 178–190.

[14] Jeffrey West et al. "A survey of open questions in adaptive therapy: Bridging mathematics and clinical translation". In: *Elife* 12 (2023), e84263.

[15] Nathan Moore, JeanMarie Houghton, and Stephen Lyle. "Slow-cycling therapy-resistant cancer cells". In: *Stem cells and development* 21.10 (2012), pp. 1822–1830.

[16] H J Broxterman et al. "Induction by verapamil of a rapid increase in ATP consumption in multidrug-resistant tumor cells." eng. In: *FASEB J* 2.7 (1988), pp. 2278–2282. ISSN: 0892-6638 (Print); 0892-6638 (Linking). DOI: 10.1096/fasebj.2.7.3350243.

[17] Maximilian AR Strobl et al. "Turnover modulates the need for a cost of resistance in adaptive therapy". In: *Cancer research* 81.4 (2021), pp. 1135–1147.

[18] Jill A Gallaher et al. "Spatial Heterogeneity and Evolutionary Dynamics Modulate Time to Recurrence in Continuous and Adaptive Cancer Therapies." eng. In: *Cancer Res* 78.8 (2018), pp. 2127–2139. ISSN: 1538-7445 (Electronic); 0008-5472 (Print); 0008-5472 (Linking). DOI: 10.1158/0008-5472.CAN-17-2649.

[19] Yannick Viossat and Robert Noble. "A theoretical analysis of tumour containment." eng. In: *Nat Ecol Evol* 5.6 (2021), pp. 826–835. ISSN: 2397-334X (Electronic); 2397-334X (Linking). DOI: 10.1038/s41559-021-01428-w.

[20] L. M. Sonneborn and F. S. Van Vleck. "The Bang-Bang Principle for Linear Control Systems". In: *Journal of The Society for Industrial and Applied Mathematics, Series A: Control* 2 (1964), pp. 151–159. URL: https://api.semanticscholar.org/CorpusID:123110370.

[21] Alfred J. Lotka. "ELEMENTS OF PHYSICAL BIOLOGY". In: *Science Progress in the Twentieth Century (1919-1933)* 21.82 (1926), pp. 341–343. ISSN: 20594941. URL: http://www.jstor.org/stable/43430362 (visited on 08/17/2023).

[22] Jessica Cunningham et al. "Optimal control to reach eco-evolutionary stability in metastatic castrate-resistant prostate cancer". In: *Plos one* 15.12 (2020), e0243386.

[23] Masud MA, Jae-Young Kim, and Eunjung Kim. "Containing Cancer with Personalized Minimum Effective Dose". In: *BioRxiv* (2022), pp. 2022–03.

[24] Wolfram Research, Inc. *Mathematica, Version 14.0*. Champaign, IL, 2024. URL: https://www.wolfram.com/mathematica.
